# Amyloidogenic proteins in the SARS-CoV and SARS-CoV-2 proteomes

**DOI:** 10.1101/2021.05.29.446267

**Authors:** Taniya Bhardwaj, Kundlik Gadhave, Shivani K Kapuganti, Prateek Kumar, Zacharias Faidon Brotzakis, Kumar Udit Saumya, Namyashree Nayak, Ankur Kumar, Neha Garg, Michele Vendruscolo, Rajanish Giri

## Abstract

The phenomenon of protein aggregation is associated with a wide range of human diseases. Our knowledge on the aggregation behaviour of viral proteins, however, is still rather limited. Here, we investigated this behaviour in the the SARS-CoV and SARS-CoV-2 proteomes. An initial analysis using a panel of sequence-based predictors suggested the presence of multiple aggregation-prone regions in these proteomes, and revealed an enhanced aggregation propensity in some SARS-CoV-2 proteins. We then studied the in vitro aggregation of predicted aggregation-prone SARS-CoV-2 proteins, including the signal sequence peptide and fusion peptide 1 of the spike protein, a peptide from the NSP6 protein (NSP6-p), the ORF10 protein, and the NSP11 protein. Our results show that these peptides and proteins form aggregates via a nucleation-dependent mechanism. Moreover, we demonstrated that the aggregates of NSP11 are toxic to mammalian cell cultures. These findings provide evidence about the aggregation of proteins in the SARS-CoV-2 proteome.

**Significance:** The aggregation of proteins is linked with human disease in a variety of ways. In the case of viral infections, one could expect that the aberrant aggregation of viral proteins may damage the host cells, and also that viral particles may trigger the misfolding and aggregation of host proteins, resulting in damage to the host organism. Here we investigate the aggregation propensity of SARS-CoV-2 proteins and show that many of them can form aggregates that are potentially cytotoxic. In perspective, these results suggest that a better understanding of the effects of viruses on the human protein homeostasis system could help future therapeutic efforts.

## Introduction

The ongoing Covid-19 pandemic, which is caused by the SARS-CoV-2 virus, has caused devastating loss of human lives and disruptions of the global economy (1, 2). Intense studies are ongoing to understand the molecular mechanisms of SARS-CoV-2 pathogenesis in order to develop effective drugs.

Given the widespread nature of the phenomenon of protein misfolding and aggregation, it is important to study its manifestation and possible implications in the case of SARS-CoV-2 proteome. Quite generally, there are at least three different aspects of the interplay between the viral and host proteomes in terms of protein aggregation: (i) the functional aggregation of viral proteins may help the virus in hijacking the replication machinery of a host cell (3, 4), (ii) the aberrant aggregation of viral proteins may represent an additional mechanism by which viruses damage the host cells (5, 6), (iii) the viral particles can trigger the misfolding and aggregation of host proteins, resulting in damage to the host organism, a process that for some viruses has been linked with the onset of Alzheimer’s disease (7).

Some viral proteins are known to form amyloid aggregates that are implicated in the viral pathogenesis. An example is that of the protein PB1 of the influenza A virus, which forms one of the three subunits of the viral polymerase. During the early stages of viral infection, PB1 is expressed in its monomeric form, but then accumulates into amyloid-like forms at a later stage of infection, which are toxic to the infected cells (5)(8, 9). Another example concerns a highly mutating H1N1 influenza A virus, whose nuclear export protein exhibits an intrinsic property to form aggregates, which has been correlated with its role in virion budding (10, 11). Viral pathogenesis mediated by viral protein aggregates has also been shown for the protein M45 of murine cytomegalovirus (6). We also note that viral capsid proteins may be prone to aberrant aggregation upon dysregulation of the functional self-assembly process, a process dependent on the environmental conditions and proceeds via nucleation and growth (12).

It has been reported that in SARS-CoV, which is closely related to SARS-CoV-2, protein M, one of the membrane-forming protein of the virus, can undergo aggregation (13). It has also been shown that the C-terminal end of protein E of SARS-CoV includes an aggregation-prone motif, and that this peptide can form aggregates in vitro (14). Furthermore, it is known that the transmembrane domain (TMD) of protein E oligomerizes to form pentameric non-selective ion channels that might act as a viroporin, a small membrane-embedded protein having ion-conducting properties (15). In SARS-CoV, aggregated forms of ORF8b induce endoplasmic reticulum stress, lysosomal damage, and activation of autophagy (16).

Following these reports about the aggregation of coronavirus proteins, here we aimed to investigate the aggregation propensity and the aggregation-prone regions (APRs) of the proteins of in the SARS-CoV-2 proteome. The RNA genome of SARS-CoV-2 encodes 29 proteins, which can be divided into three classes: structural, accessory, and non-structural proteins (**Figure 1**) (17). Our results identify non-structural proteins (NSPs), which play a crucial role during the initial and transitional phases of the viral life cycle in the host cell, as particularly aggregation prone. We also compared the aggregation propensities of the SARS-CoV-2 proteins with those of the SARS-CoV proteins. We then focused our analysis on specific SARS-CoV-2 proteins by further investigating the signal sequence peptide and fusion peptide 1 of the spike protein, full-length ORF10, NSP6-p (residues 91-112 of NPS6), and NSP11 using fluorescence and spectroscopy methods, as well as atomic force microscopy (AFM) to visualize the morphology of resultant aggregates. In addition, we investigated the cytoxicity of NSP11 aggregates on different mammalian cell lines.

**Figure 1.**
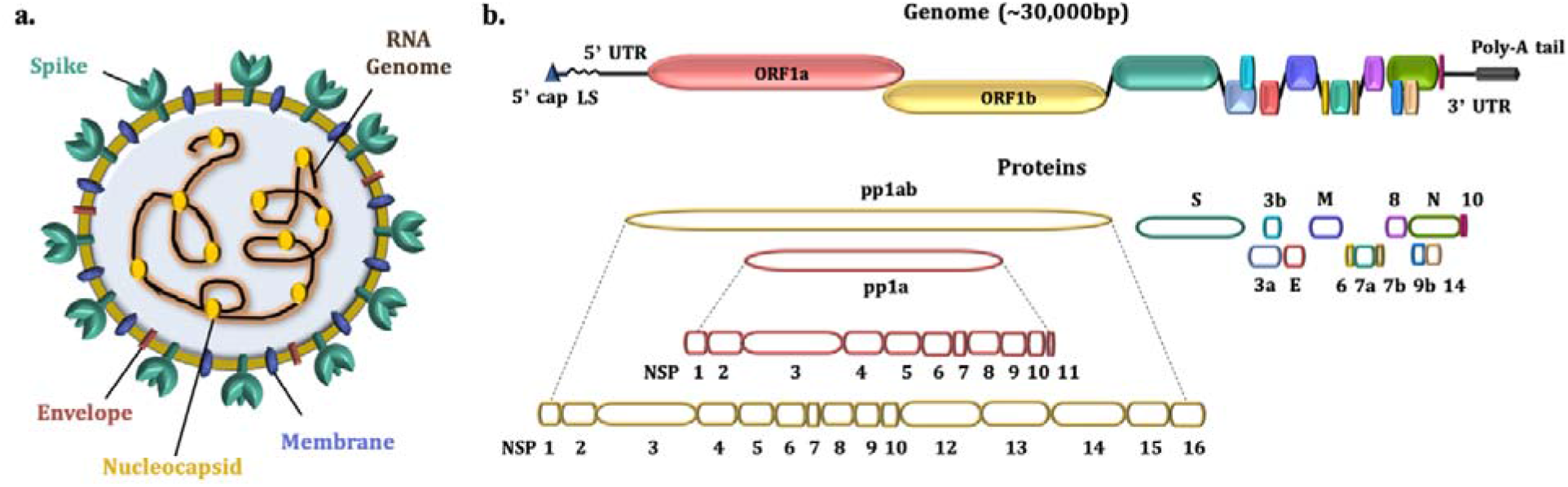
Schematic illustration of the organisation of the genome and proteome of SARS-CoV-2. (**a**) The SARS-CoV-2 viral particle comprises positive-sense single-stranded RNA, which is associated with the nucleocapsid protein (N), and three surface proteins, spike (S), membrane (M), and envelope (E), which are embedded in the lipid bilayer. (**b**) The length of the SARS-CoV-2 genome is ~30 kbp, which encodes for four structural, nine accessory, and sixteen non-structural proteins.

## Results

### Aggregation-prone regions in the SARS-CoV and SARS-CoV-2 proteome

The tendency to self-assemble into amyloid structures is an intrinsic property of proteins and depends on the presence of APRs within their amino acid sequences (18, 19). This tendency is in competition with that of self-assembling to form functional complexes (20). To investigate this competition, in this study we analyzed the tendency of SARS-CoV and SARS-CoV-2 proteins to form amyloid aggregates using different computational prediction methods. We employed a combination of three different individual predictors - FISH Amyloid, AGGRESCAN, and FoldAmyloid, and a meta-predictor MetAmyl to analyse the presence of aggregation-prone regions. We also used CamSol to predict the solubility of the proteins of both viruses. APRs predicted by different servers are abundant in both proteomes, and all proteins were found to contain at least one APR (**Tables S1-S6**).

For the comparison of amyloidogenic propensity of the SARS-CoV-2 and SARS-CoV proteomes, we calculated the the mean predicted percentage amyloidogenic propensity (PPAP) from the aggregation-prone regions obtained from the four predictors (MetAmyl, FISH Amyloid, AGGRESCAN, and FoldAmyloid) for both viruses (**Figure 2a,b**). Numerous proteins, particularly accessory proteins of SARS-CoV-2 (**Figure 2a**) were observed to be more amyloidogenic as compared with accessory proteins of SARS-CoV (**Figure 2b**).

**Figure 2.**
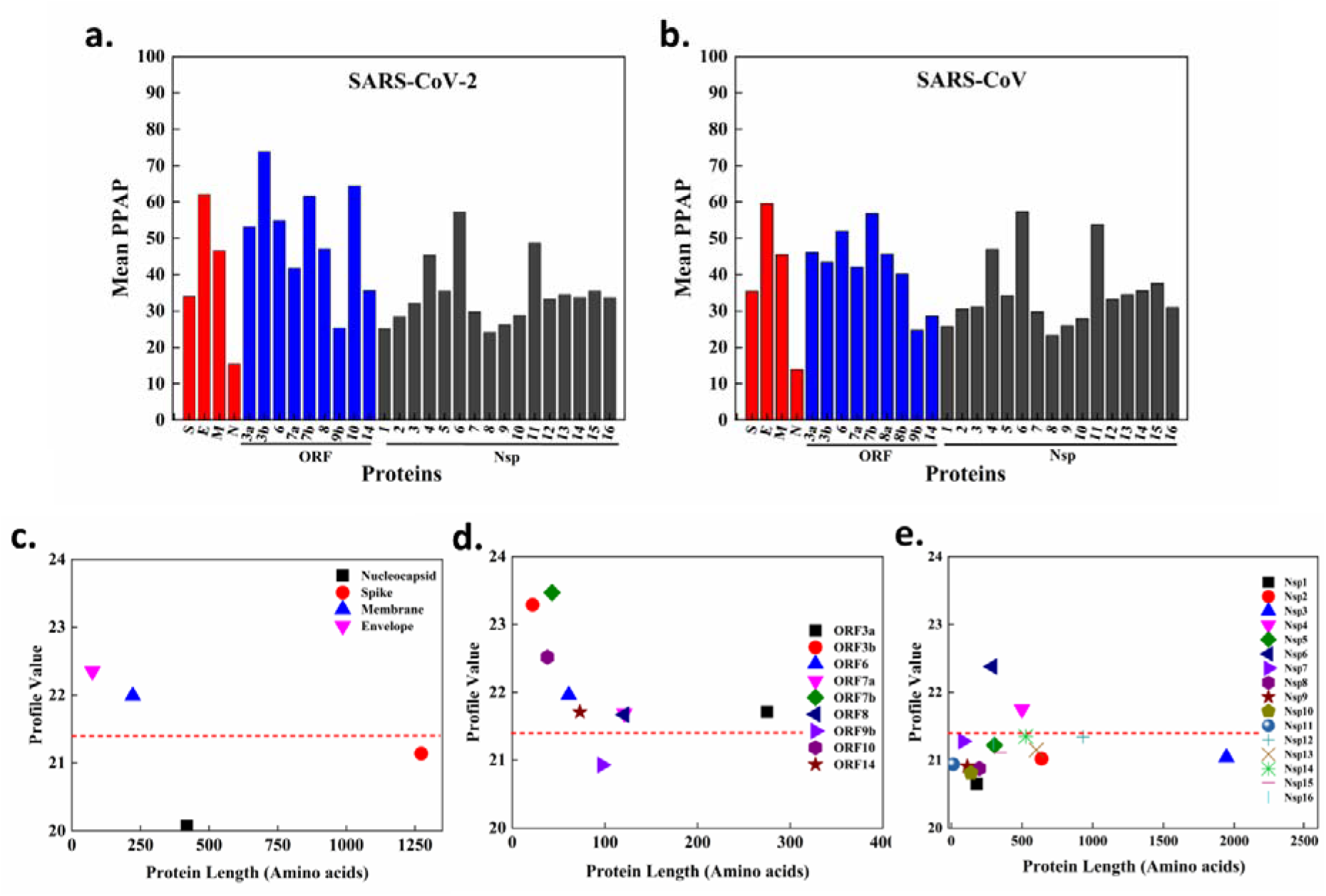
Amyloidogenic propensity analysis of the SARS-CoV and SARS-CoV-2 proteomes. **(a,b)** Mean predicted percentage amyloidogenic propensity (PPAP) calculated using the mean of percentage of aggregation prone regions obtained from four servers (MetAmyl, AGGRESCAN, FoldAmyloid, FISH Amyloid) for SARS-CoV-2 **(a)** and SARS-CoV **(b)**. **(c-e)** Average profile value for protein obtained from FoldAmyloid analysis of proteins against protein length for structural proteins **(c)**, accessory proteins **(d)**, and non-structural proteins **(e)**. The analysis was done at default settings in the FoldAmyloid server (threshold: 21.4, represented by the red-colored short-dashed line, and scale, i.e. the expected number of contacts within 8 Å).

In SARS-CoV-2, among the structural proteins, M and E are found to be more amyloidogenic by FoldAmyloid in comparison with N and S (**Figure 2c and Table S1**), and the accessory proteins to have more abundance of APRs (**Table S2**). Except for ORF9b, all other proteins are showing FoldAmyloid profile value above the threshold value (**Figure 2d and Table S3**). These 16 NSPs perform diverse roles, such as evading the host immune system, protection from host defence mechanisms, virus replication, and spreading of the infection (17). From the FoldAmyloid analysis, after plotting the average profile value for protein (**Figure 2e**), NSP4 and NSP6 show profile values above the cut-off, which indicates a highly amyloidogenic nature.

To gain insights into the possible cleavage of APRs, 20S proteasome cleavage sites within the entire SARS-CoV-2 proteome were predicted by the NetChop 3.1 server (**Tables S7-S9**). The reason of identifying these sites, in context of our study, is twofold. First, the predicted sites residing inside APRs suggest that due to aggregation the proteasome may not be able to successfully cleave the viral proteins. Secondly, the proteasome could cleave the viral proteins and release amyloidogenic reagions inside the host cell. Here, in case of accessory proteins, we found many cleavage sites located in APRs (**Tables S7-S9**).

### Prediction of aggregation-prone regions in the structural proteins

Four structural proteins of SARS-CoV-2 participate in the virion assembly and packaging processes and in providing structure to the virus (23). We analyzed the APRs of these structural proteins (**Table S1**). One of these proteins, S, is a heavily glycosylated transmembrane protein whose N-terminal S1 domain harbours receptor binding sites for the host cell, and C-terminal S2 domain mediates the fusion between virion and host cells (24). We predicted several APRs in S (**Table S1**).

We note that a recent cryo-EM study revealed a hinge motion of S1, which take the virus from its active state to its inactive state (25). To investigate a possible relationship between Covid-19 and Alzheimer’s disease (7), we performed a multiple sequence alignment (MSA) of the Aβ42 peptide with S (**Figure S1a**), observing that Aβ42 containes a similar amyloido-genic region (_26_SNKGAI_31_) as detected in S (_968_SNFGAI_973_). By inspecting frames in the open state (**Figure S1b**), we noted that this APR can be exposed in the active state, thereby suggesting a possible mechanism by which the virus could influence the aggregation of Aβ42.

We also predicted several APRs in the other structural proteins (M, E and N) (**Table S1**). The membrane protein M gives shape to the virus, promotes viral membrane curvature, and binds with the nucleocapsid RNA complex during virus packaging. The envelope protein E is a transmembrane protein with an ion channel activity that facilitates the assembly and release of viral particles. The amyloid-forming propensity of 9-residue stretch (VYVYSRVK) of E have been reported previously for SARS-CoV (14). The nucleocapsid protein N is the proteinaceous part of the viral nucleocapsid, which interacts with viral RNA and helps its packaging into the virion. According to our analysis, AGGRESCAN detected a total of 18% amyloidogenic regions in SARS-CoV-2 N and only 16% in SARS-CoV N. Similar to these results, FoldAmyloid, also predicted only 11% amyloidogenic regions in SARS-CoV N, ~3% less than SARS-CoV-2 N, which contains a total of 14% amyloidogenic regions.

### Prediction of aggregation-prone regions in the accessory proteins

The coronavirus genome codes for proteins termed accessory, which are multifunctional proteins that play an important role in modulating the host response to virus infection, such as down-regulation of interferon pathways, the release of proinflammatory cytokines and chemokines, and the induction of autophagy. According to predictors used in this study, all accessory proteins have multiple aggregation-prone regions (**Table S2**). ORF3a, and the N-terminal regions of ORF6, ORF7a, ORF7b, ORF8, and ORF10 are found to have a tendency for aggregation. In addition, the C-terminal regions of ORF7a, ORF8, ORF9b, ORF10, and ORF14 proteins also exhibit numerous APRs.

### Prediction of aggregation-prone regions in the non-structural proteins

The SARS-CoV-2 proteome includes 16 non-structural proteins, all of which were found to comprise APRs in their sequences (**Table S3**), which could play a role in their behaviours and interactions with the host cells. NSP1 is involved in creating a suitable environment for viral propagation by blocking host cell translation and host immune response. NSP3 is a papain-like protease that cleaves the viral ORF1ab polyprotein at the NSP1/NSP2, NSP2/NSP3, and NSP3/NSP4 boundaries, and interferes with the proper functioning of the host proteome by blocking the cellular degradation system. NSP4 and NSP6 are transmembrane proteins that may have a scaffolding activity for viral replication vesicles formation (26). NSP5 is a serine like protease that catalyzes the rest eleven cleavage events of the ORF1ab polyprotein. NSP12 is an RNA-dependent RNA polymerase, and NSP7 and NSP8 function as its processivity clamps. NSP10 is a cofactor for NSP16, which protects viral RNA from host antiviral measures. NSP13 is an RNA helicase and NSP14 is a methyltransferase that adds a 5’ cap to viral RNA and is involved in proofreading of the viral genome by virtue of its has 3’-5’ exonuclease activity. NSP15 is an endoribonuclease that has a defensive role from host attacks.

### Experimental analysis of aggregation-prone SARS-CoV-2 proteins

After the computational prediction of APRs in the SARS-CoV and SARS-CoV-2 proteomes, we investigated the *in vitro* aggregation behaviour of various SARS-CoV-2 proteins and peptides. For this purpose, we selected pH 7.4 and temperature 37 °C, and traced the aggregation process using the fluorescent dye thioflavin T (ThT), which interacts with amyloid fibrils and gives a maximum emission peak at ~490 nm upon binding β-sheets in amyloid fibrils (27)(28). As protein aggregation often occurs via a nucleation-polymerization mechanism (29, 30), we studied this reaction using ThT fluorescence (λ_max_ at 490 nm) in presence of a fixed volume of incubated samples (25 μM).

Then, we employed AFM to gain insights about the morphological features of the aggregates. To this end, we studied the *in vitro* aggregation of the spike signal peptide and fusion peptide 1, ORF10 protein, NSP6-p, and NSP11 proteins of SARS-CoV-2. As a negative control, we studied the peptide corresponding to residues 131-180 of NSP1 (NSP1-p), which is not predicted to contain APRs by any of the software used, finding that it does not form amyloid-like aggregates and it does not show any change in the ThT assay.

### Spike signal peptide

Spike plays a key role in the receptor recognition and cell membrane fusion process. In its mature form, spike contains four regions, a signal sequence, ectodomain, transmembrane domain and endodomain, which is further divided into the S1 and S2 subunits. The 12-residue N-terminal signal sequence (SP) directs spike to its destination in the viral membrane (31). CamSol predicted SP to be poorly soluble and all aggregation prediction servers identified it as an APR (**Table S1**). Our study of the aggregation behaviour of SP *in vitro* (see Methods) showed an over three-fold increase in ThT fluorescence intensity upon aggregation (**Figure 3a**). The nucleation process exhibited little to no lag phase and reached saturation after 10 hours with an half-time (T_1/2_) of about 4 hours (**Figure 3b**). We then used atomic force microscopy (AFM) to gain more information and confirm the changes in structural features upon aggregation. AFM images of SP aggregates displayed short but slender fibrils formed after 170 hours (**Figure 3c-f**).

**Figure 3.**
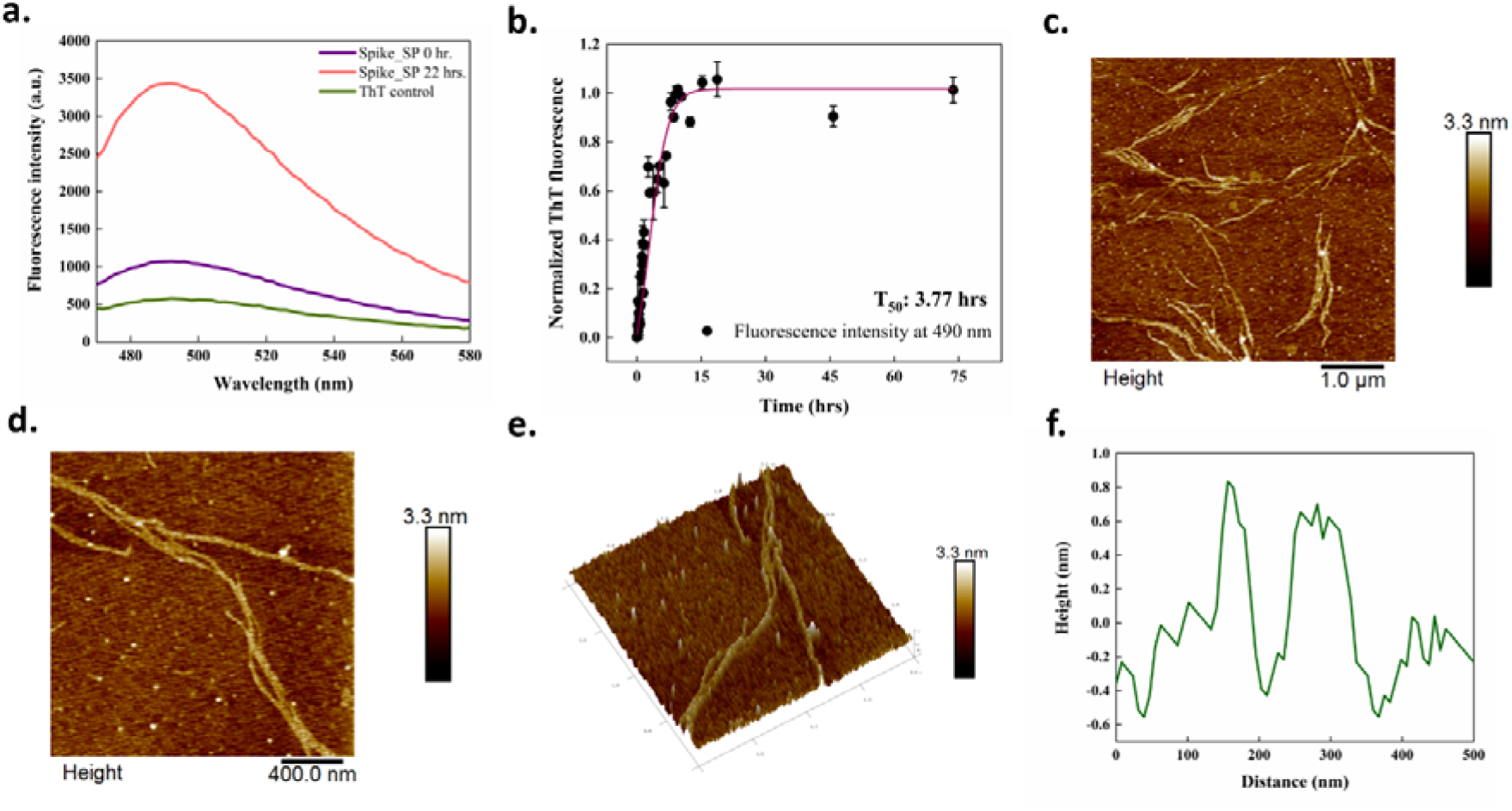
In vitro aggregation of the signal peptide (SP) of the SARS-CoV-2 spike protein. **(a)** ThT fluorescence scan at 440 nm excitation and 470-700 nm emission. (**b**) ThT fluorescence kinetics of the SP. Black circles represent ThT fluorescence intensity at 490 nm, and the line represents sigmoidal fitting. (**c-e)** Morphology of aggregates after a 170 hour incubation. Scale bars represent 1 μm and 400 nm for zoomed image. (**f**) Height profile of SP fibrils from a zoomed image of panel **e**.

### Spike fusion peptide 1

Spike contains a fusion peptide (FP1) of 15-20 residues in the S2 subunit that helps the virus penetrate the host cell membrane (31, 32). All the predictors used in this study identified APRs in this region. We thus analysed the aggregation of FP1 in vitro finding an increment in ThT fluorescence intensity (**Figure 4a**). Upon a 3.5 hour incubation, we found an increased ThT fluorescence by ~6-fold in comparison to a freshly dissolved peptide sample. Using ThT fluorescence at 490 nm, the aggregation of FP1 reached saturation in about 3 hours with a T_1/2_ of about 1 hour (**Figure 4b**). Tapping-mode AFM used to investigate the morphological features of aggregates in this study exposed the presence of numerous entangled filaments (**Figure 4c,d**). These differently sized filamentous FP1 aggregates at 96 hours have a height distribution peaking at ~6 nm (**Figure 4e**). For FP1, the ThT fluorescence readings were maintained at saturation while preparing the samples for AFM imaging.

**Figure 4.**
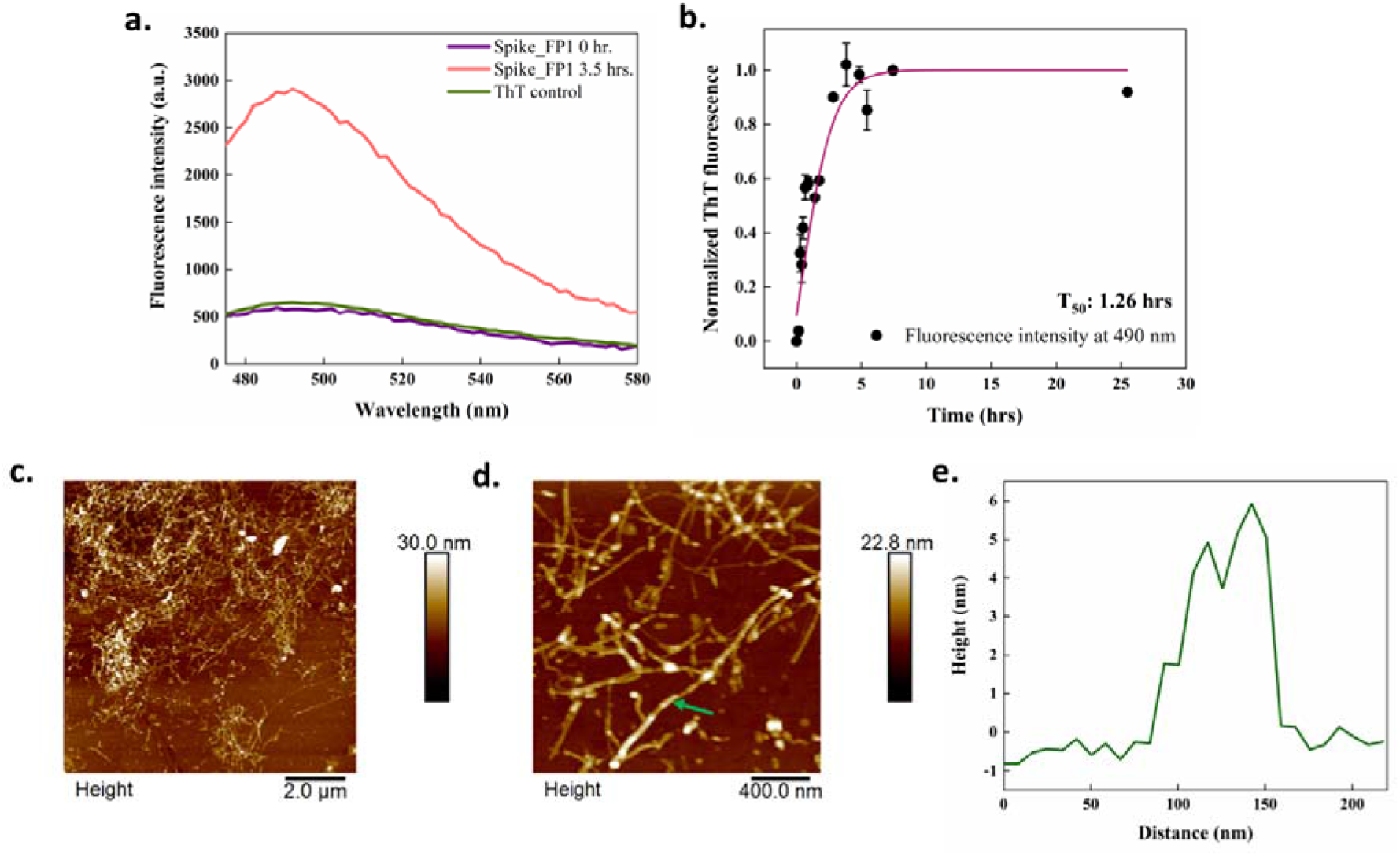
In vitro aggregation of fusion peptide 1 (FP1) of the SARS-CoV-2 spike protein. **(a)** ThT fluorescence scan at 440 nm excitation and 470-700 nm emission scan. (**b**) ThT fluorescence kinetics of FP1. Black circles represent ThT fluorescence intensity at 490 nm, and the line represents a sigmoidal fitting. **(c**,**d)** Morphology of FP1 aggregates after a 96-hour incubation. Scale bars represent 2 μm and 400 nm for the zoomed image. (**e**) Height profile of FP1 fibrils shown with a green arrow from the zoomed image in panel **d**.

### ORF10 protein (full-length)

The SARS-CoV-2 ORF10 protein is predicted to be translated into a 38-residue long protein that does not have significant homology with any known proteins. Although the evidence of presence of ORF10 in SARS-CoV-2 infected cells is limited (33, 34), it has been investigated to have a high dN/dS value (nonsynonymous over synonymous substitution rate) (3.82) (35). We predicted ORF10 to contain multiple APRs (**Table S2**). In *in vitro* aggregation experiments, ORF10 showed a ~7-fold increase in ThT fluorescence after 11 days of constant stirring at 1000 rpm and incubation at 37 °C (**Figure 5a**). The T_1/2_ of aggregation kinetics of this process was found to be ~30 hours (**Figure 5b**). In comparion with fibrillar aggregates of other peptides studied in this report, ORF10 aggregates were visualized to form amorphous aggregates using AFM (**Figure 5c-e**). The ThT fluorescence of ORF10 protein aggregates were maintained at saturation while preparing the samples for AFM imaging.

**Figure 5.**
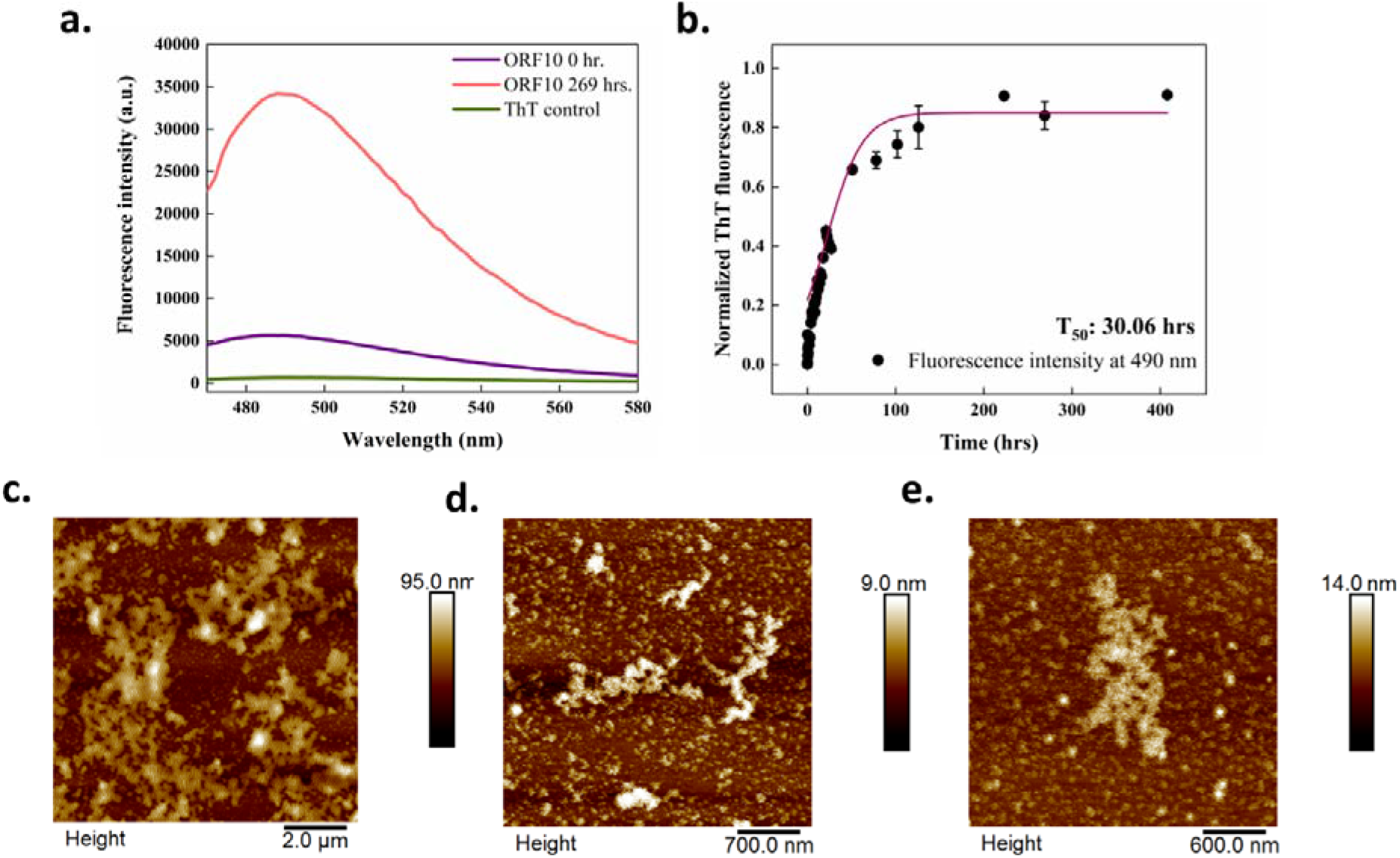
In vitro aggregation of the ORF10 protein of SARS-CoV-2. **(a)** ThT fluorescence scan at 440 nm excitation and 470-700 nm emission. (**b**) ThT fluorescence kinetics of ORF10 protein; black circles represent the ThT fluorescence intensity at 490 nm, and the line represents a sigmoidal fitting. (**c**-**e**) Morphology of aggregates after a 96-hour incubation; scale bars represent 2 μm, 700 nm and 600 nm for AFM images illustrated in **c**, **d**, and **e** respectively.

### NSP6-p

The SARS-CoV-2 NSP6 protein is one of an essential host immune system antagonist. It antagonizes IFN-I signaling through blocking TANK binding kinase 1 (TBK1) to suppress interferon regulatory factor 3 (IRF3) phosphorylation efficiently than the SARS-CoV and MERS-CoV NSP6 proteins (36). Whilst accomodating multiple transmembrane regions, the region of residues 91-112 of NSP6 is of particular importance since it lies outside the membrane and thereby can interact with host proteins (37). This region exhibited APRs using all predictors (**Table S3**). The intrinsic solubility of this region is also calculated to low by CamSol server. As an experimental validation, we studied that aggregation of the NSP6-p peptide (residues 91-112 of NSP6) in buffered conditions. Its aggregation using ThT fluorescence is detected at ~40 hours (**Figure 6a**). A kinetic analysis revealed a ~15 hours long lag phase followed by an exponential phase attainting the final plateau phase (**Figure 6b)**. The T_1/2_ value of aggregation kinetics of NSP6-p was calculated to be about 21 hours. **Figure 6c-e** reports the AFM images of a 96-hour aggregated sample. The height profile shows the peak height of ~3 nm of an aggregated fibril **(Figure 6f)**.

**Figure 6.**
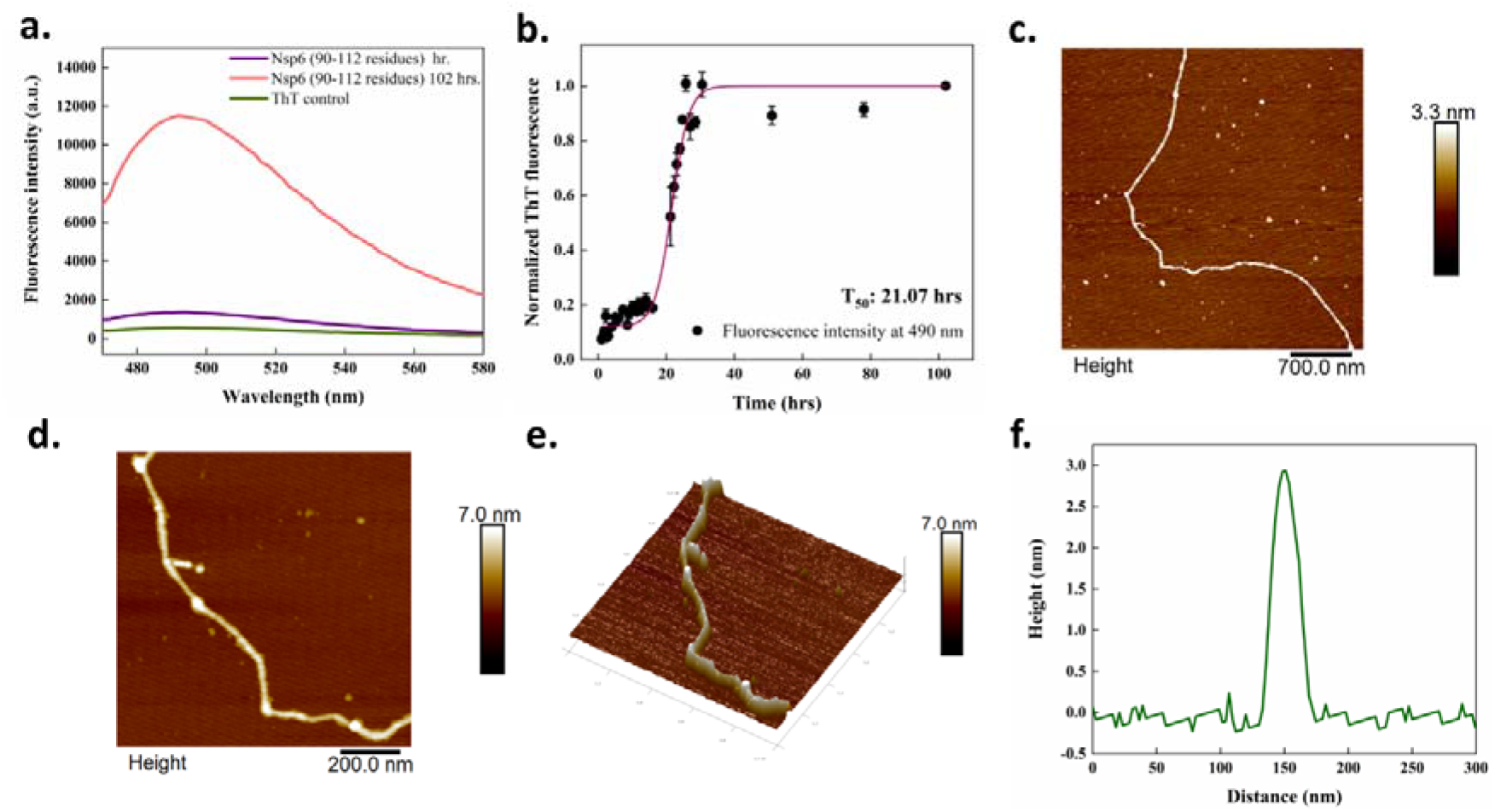
In vitro aggregation of NSP6-p. **(a)** ThT fluorescence scan at 440 nm excitation and 470-700 nm emission. (**b**) ThT fluorescence kinetics of NSP6-p; black circles represent ThT fluorescence intensity at 490 nm, and the line represents a sigmoidal fitting. **(c-e**) Morphology of NSP6-p fibrils after a 96-hour incubation; scale bars represent 200 nm and 100 nm for zoomed image in 2D. (**f**) Height profile of NSP6-p fibrils from a zoomed image of panel **e**.

### NSP11 protein

NSP11 of SARS-CoV-2 is a 13 amino acid length peptide, cleaved from polyprotein 1a. The first 9 residues of NSP11 are similar to the first 9 residues of NSP12 (RdRp) (38). For the identification of amyloid-forming residues and the aggregation propensity of NSP11, we used four aggregation prediction servers. AGGRESCAN predicted the region 6-13, FISH Amyloid predicted 6-10, and MetAmyl Predicted 8-13 in NSP11 as APRs (**Table S3**). ThT dye binds to NSP11 amyloid fibrils (192-hour incubated sample) and increases its fluorescence intensity as compared to the freshly-dissolved NSP11 monomers (**Figure 7a**). Futhermore, according to a kinetic analysis of NSP11 aggregation, following a lag phase of ~45 hours, the process reached a plateau phase after 110 hours (**Figure 7b)** with T_1/2_ of 64 hours. After incubating NSP11 for 192 hours, we detected typical amyloid fibrils using AFM (192 h) of ~2.5-3 nm height (**Figure 7d,e**).

**Figure 7.**
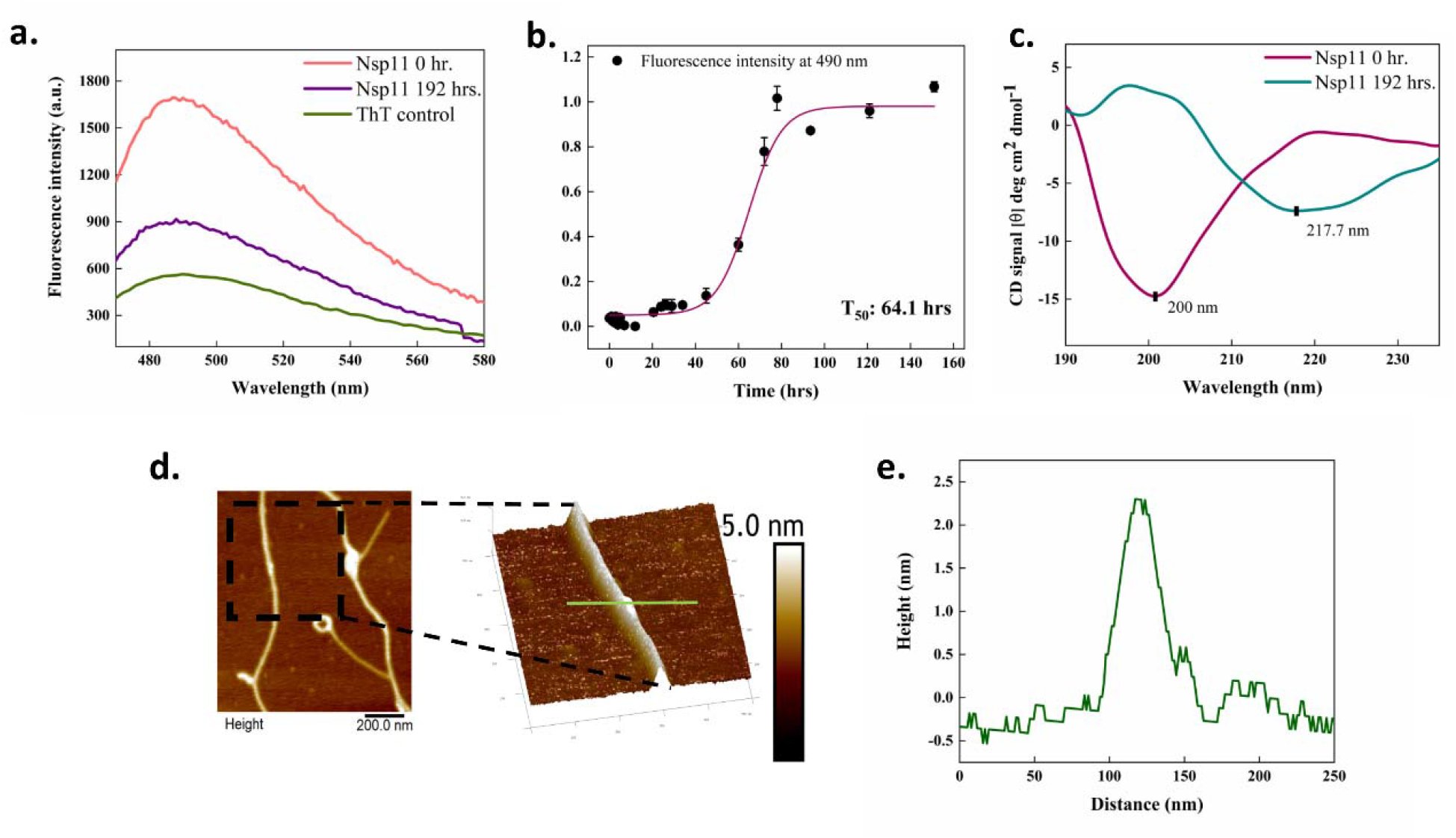
In vitro aggregation of the SARS-CoV-2 NSP11 protein. (**a**) ThT fluorescence scan at 440 nm excitation and 470-700 nm emission. (**b**) ThT fluorescence kinetics of NSP11; black circles represent ThT fluorescence intensity at 490 nm, and the line represents a sigmoidal fitting. **(c)** CD spectra of NSP11 0-hour (monomer) and 192-hour (aggregate) samples. (**d**) Morphology of NSP11 fibrils after 192 hours incubation obtained using AFM. (**e**) AFM height profile of NSP11 fibril from a zoomed image of panel **d**.

We then monitored by far-UV CD-spectroscopy the changes in the secondary structure during the aggregation of NSP11. The far-UV CD spectrum of monomeric NSP11 is representative of the disordered proteins, which is also reported in our previous study (39), represents a robust negative band near 200 nm (**Figure 7c**). However, after incubating NSP11 at 37 °C in phosphate buffer (20 mM phosphate, 50 mM NaCl, pH 7.4) at 1000 rpm for 192 hours, we observed a gain of weak negative band, appearing as a shoulder around 218 nm, suggesting the presence of β-sheet secondary structure elements in aggregated NSP11 protein (**Figure 7c**). These results indicate that, upon aggregation, NSP11 converts from disordered to a conformation with a considerable secondary structure.

### Toxic effects of NSP11 aggregates on the viability of mammalian cells

Since studies on the amyloid aggregation of viral proteins may reveal an additional potential of the viruses to damage host cells, we investigated whether or not β-sheet rich amyloid fibrils of SARS-CoV-2 NSP11 could be cytotoxic to human cells. We performed MTT assay, a colorimetric-based cell viability test to assess the effect of aggregates on the SH-SY5Y and HepG2 cell lines. The cells were treated with varying concentrations of NSP11 monomers (used as control) and amyloid aggregates (192 hours) separately. No significant cell death was observed after 24 hours (**Figure 8a,c**). However, when treatment was extended to 72 hours, cell death observed at the highest aggregate concentration is comparatively enhanced (**Figure 8b,d**).

**Figure 8.**
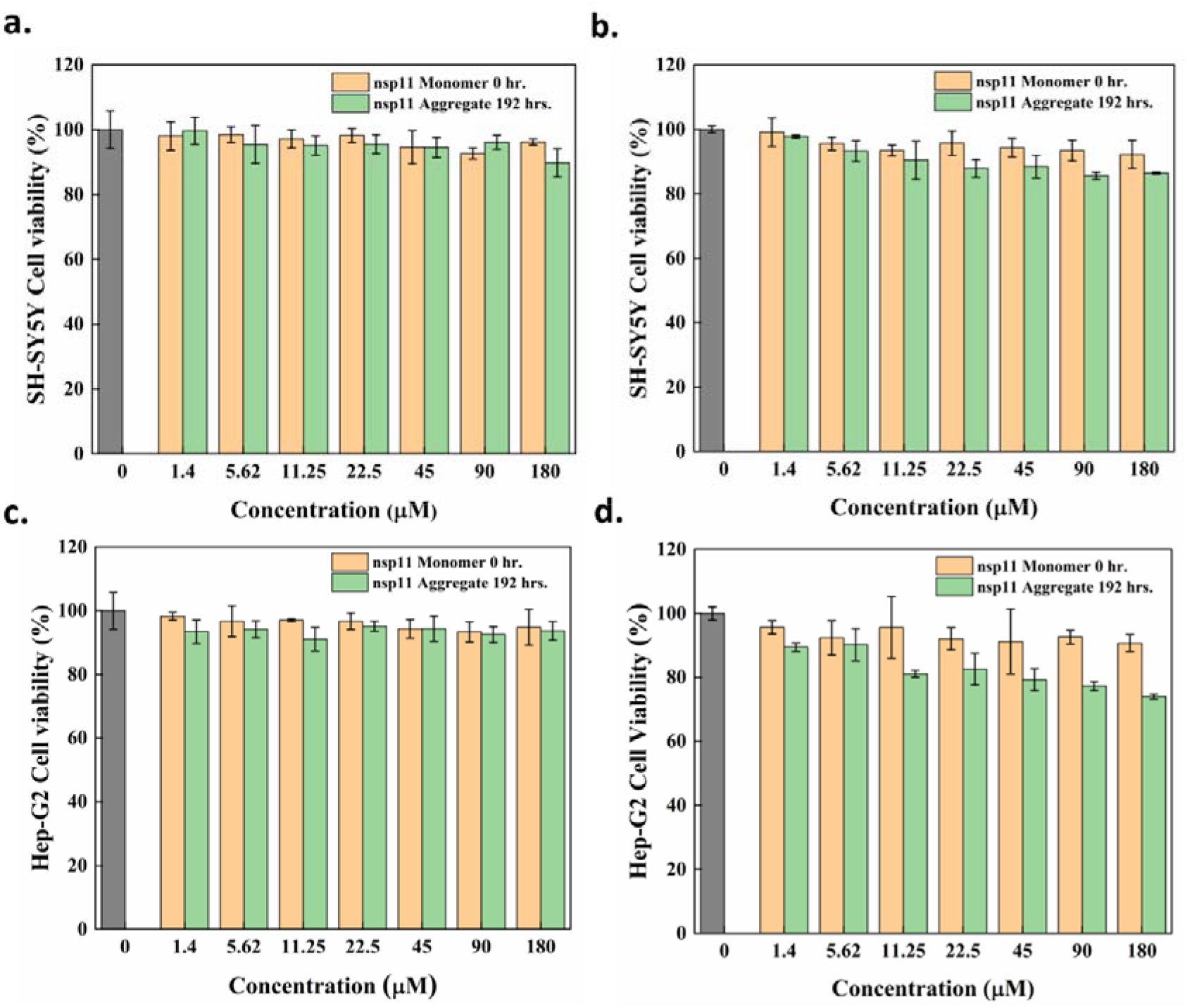
Cell viability upon NSP11 aggregation using an MTT assay. SH-SY5Y cells treated with NSP11 monomers (0-hour incubation) and amyloid fibrils (192-hour incubation) for 24 hours (**a**) and 72 hours (**b**). Hep-G2 cells were treated with NSP11 monomers (0-hour incubation) and amyloid fibrils (192-hour incubation) for 24 hours (**c**) and 72 hours (**d**). The cells treated in media containing 20 mM sodium phosphate buffer (pH 7.4) were used as a control (gray bars). The dark yellow and light green bars indicate cells treated with NSP11 monomers and fibrils, respectively; error bars represent the SEM of 3 independent samples.

We also observed that percent cell viability in HepG2 cells is reduced more than that in SH-SY5Y cells, suggesting the aggregates are comparatively more toxic to the liver cell line. HepG2 cell viability remained around 74%, while in SH-SY5Y percent cell viability remained around 86%. These results thus suggest that NSP11 aggregates are cytotoxic at relatively higher concentrations.

## Discussion and Conclusions

This study was inspired by a series of reports that associated viral infections with protein misfolding diseases of the central nervous system (CNS), including Alzheimer’s disease and related dementias (40). In particular, the Ljuangan virus was detected in the hippocampus of Alzheimer’s disease patients (41), and HIV was found to affect the levels of Aβ42 in the cerebrospinal fluid (42). An epidemiological study recently linked herpes infections with an increase in the risk of dementia (43), consistently with reports in which HSV-2 was associated with a decline in cognitive abilities of aged individuals (44), as well as with other reports in which HSV-1 was linked to Alzheimer’s disease through the facilitatation of the formation of amyloid-like structures in neural stem cells and bioengineered brain tissue cultures (45). Furthermore, the H5N1 influenza virus was reported to induce acute neurological signs such as motor disturbances and mild encephalitis in animal models resulting in CNS disorders of protein aggregation like Alzheimer’s and Parkinson’s diseases (46).

Among coronaviruses, SARS-CoV has been reported to target the brain in mice (47) and was isolated from brain tissues of a Covid-19 patient with severe CNS symptoms (48). Another case study showed the development of polyneuropathy in SARS-infected patients after the first outbreak in 2004 (49). It was then reported that SARS-CoV can enter the brain via the olfactory bulb, and that intracranial inoculation can cause extensive neuronal infection leading to death (50). Likewise, SARS-CoV-2 was associated with severe neurological manifestations, including acute cerebrovascular diseases, skeletal muscle injury, impaired consciousness, and acute hemorrhagic necrotizing encephalopathy (51, 52). The presence of SARS-CoV-2 in the cerebrospinal fluid of patients has been linked with meningitis and meningoencephalitis (53, 54), and neurons infected with SARS-CoV-2 displayed altered distribution of the tau protein (55).

To increase our understanding of the links between Covid-19 infection and pathological events linked to protein aggregation, in this study we have investigated several proteins of SARS-CoV and SARS-CoV-2 and shown that they exhibit an intrinsic propensity to aggregate. We have then demonstrated that in SARS-CoV-2 the signal peptide and fusion peptide 1 of spike, NSP6-p, ORF10, and NSP11 aggregate under in vitro conditions. Furthermore, we have also identified the cytotoxic effects of NSP11 aggregates in SH-SY5Y and HepG2 cells.

In conclusion, in this study we have analyzed the aggregation propensity of the SARS-CoV and SARS-CoV-2 proteomes and identified amyloidogenic proteins with diverse functions in the viral pathogenesis. Based on these predictions, we have demonstrated that the SARS-CoV-2 spike signal sequence peptide and fusion peptide 1, ORF10, NSP6-p, and NSP11 can form amyloid aggregates. We have then shown that NSP11 aggregates can be toxic to mammalian cells. Taken together, these results provide evidence of the tendency of several proteins in the SARS-CoV-2 proteome to form aggregates.

## Materials and Methods

### Materials

#### Peptides

The following proteins and peptides are chemically synthesized from GenScript, USA: SARS-CoV-2 Spike signal sequence (SP, residues 1-12, 92.6% purity), SARS-CoV-2 Spike fusion peptide (FP1, residues 816-837, 79% purity), SARS-CoV-2 NSP6-p (88.5% purity), SARS-CoV-2 NSP11 protein (residues 4393-4405, > 72.9% purity), NSP1-p (82% purity). SARS-CoV-2 ORF10 protein (full lenth, crude, Mass spectrometry data of crude peptides is included in supplementary information) was purchased from Thermo Scientific, USA.

#### Chemicals

TEM Grids and MICA sheets were obtained from TED PELLA INC., USA. Other chemicals, including ammonium molybdate, thioflavin T (ThT), and 3-(4,5-dimethylthiazol-2-yl)-2,5-diphenyltetrazolium bromide (MTT) were procured from Sigma Aldrich, St. Louis, USA. The chemicals used in the cell culture study were purchased from Gibco^™^. The cell lines (SH-SY5Y neuroblastoma and Hep-G2) were obtained from the National Centre for Cell Sciences (NCCS), Pune, India.

### Prediction of amyloidogenic regions in SARS-CoV-2 and SARS-CoV proteome

Aggregation-prone regions (APRs) in the SARS-CoV-2 and SARS-CoV proteome were predicted by four sequence-based methods. MetAmyl, AGGRESCAN, FoldAmyloid, and FISH Amyloid. AGGRESCAN is based on a scale for natural amino acids derived from in vivo experiments. It also assumes that short and specific sequences within the protein can regulate protein aggregation. It gives a hot spot area (HSA) score for susceptible aggregate forming residues (56). FoldAmyloid calculates the probability of backbone-backbone hydrogen bond formation and efficiently classifies the amyloidogenic peptides. It determines the amyloidogenic residues scoring above 21.4, a threshold assumed by the server (57). MetAmyl is a metapredictor and combines the strength of different individual predictors – PAFIG, SALSA, Waltz and FoldAmyloid. It creates a logistic regression model and gives score which is interpreted as probability for a fragment to form an amyloid fibril (58). FISH Amyloid is a new machine learning prediction method based on the presence of a segment with the highest scoring for co-occurrence of residue pairs (59). Additionally, CamSol is used to predict the hydrophilic and hydrophobic regions of proteins. It calculates the intrinsic solubility of proteins, which is inversely related to their aggregation propensity. CamSol assigns values to each amino acid, negative values below −1 represent insoluble residues, and positive values above +1 represent soluble residues (60). The region in a protein that is predicted with five or more than five amino acids long were considered as potentially amyloidogenic regions. Furthermore, in silico mapping of 20S proteasome cleavage sites across SARS-CoV2 proteome predicted with proteasome prediction server NetChop 3.1 algorithm, webserver tool.

### Preparation of the samples for the aggregation assays

Peptides were dissolved in a 100% HFIP to remove pre-existing aggregates, and left to evaporate at room temperature overnight resulting in dry peptides. These peptide films were then dissolved according to their hydrophobic character and solvent recommended by GenScript and Thermo Scientific, USA, before incubation for aggregation experiments, as detailed in **Table 1**. Monomeric samples for the aggregation assays were then incubated at 37 °C with constant stirring (1000 rpm) on Eppendorf ThermoMixer C.

**Table 1.**
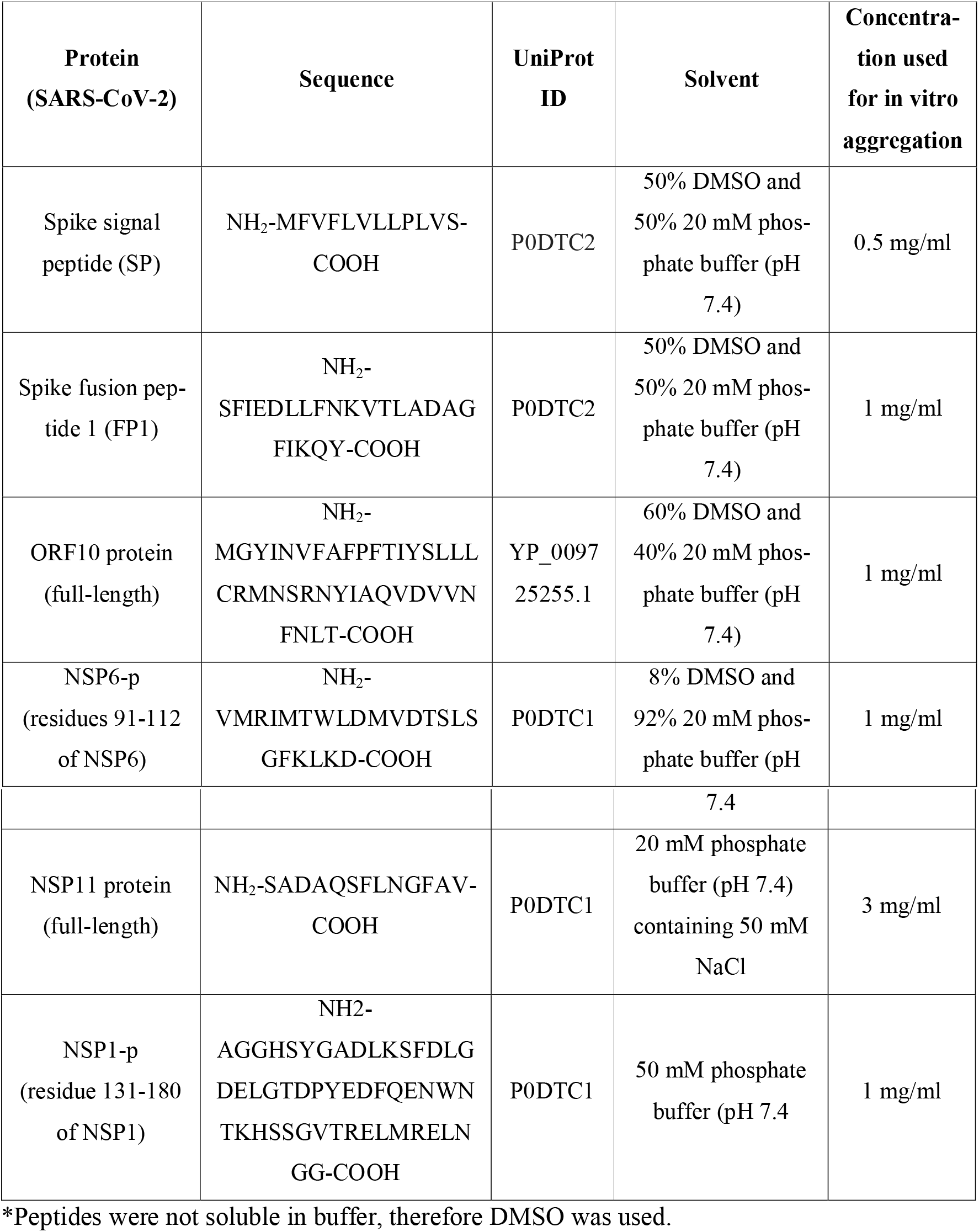
Conditions used to prepare the peptides and proteins for the in vitro aggregation assay.

### Thioflavin T aggregation assay

To analyze the aggregation process *in vitro* we used thioflavin T (ThT), which is an extensively used dye that upon binding to amyloid fibrils gives an increase in fluorescence (61, 62). 25 μM aggregated samples were prepared in 20 mM sodium phosphate buffer (pH 7.4) with 25 μM ThT and incubated for 5 minutes in dark conditions. The samples were excitated at 440 nm wavelength and emission spectra were recorded from 460 to 600 nm wavelengths using TECAN Infinite M200 PRO multimode microplate reader. Further, the kinetics of peptide aggregation were monitored at different time intervals. At different incubation times, 25 μM of peptide samples and 25 μM ThT were mixed, incubated, and fluorescence intensity of ThT was monitored at 440 nm excitation and 490 nm emission wavelength using black 96 well plates in TECAN Infinite M200 PRO multimode microplate reader. For DMSO soluble peptides, ThT dye mixed with DMSO (at particular concentration) and buffer was used as ThT control. All the measurements were set up in duplicates, and the average value was reported with standard error (SE). Data were fitted using a sigmoidal curve to obtain *T*_1/2_ (the time at which ThT fluorescence intensity reaches 50% of its maximum value)

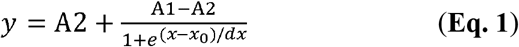

where A1 indicates the initial fluorescence, A2 the final fluorescence, *x*_0_ the midpoint (T_1/2_) value), and *dx* is a time constant. The lag time was calculated as

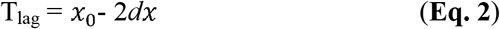

### Circular dichroism (CD) spectroscopy

Far-UV CD spectra were recorded to characterize the secondary structure of aggregated proteins and peptides. 20 mM sodium phosphate buffer (pH 7.4) containing 50 mM NaCl was used to prepare the samples, and far-UV CD spectra were recorded in 1 mm path length quartz cuvette using J-1000 series JASCO spectrophotometer system at 25 °C. The spectra were recorded from 190 nm to 240 nm wavelength with a bandwidth of 0.5 mm as an average of 3 consecutive scans. The spectrum of the buffer alone was subtracted from the spectrum of samples.

### Atomic force microscopy (AFM)

We obtained AFM images of aggregated fibrils using tapping-mode AFM (Dimension Icon from Bruker). We carried out the measurements by depositing a twenty-fold diluted aggregated solution on a freshly cleaved mica surface for each of the peptides. After incubating for 1 h, the surfaces were rinsed three times with deionized water, dried at room temperature overnight, and finally, images were recorded.

### Cell viability assays

The effect of SARS-CoV-2 NSP11 fibrils on cell viability was assessed in two different cell lines (SH-SY5Y neuroblastoma cells & HepG2 hepatocellular carcinoma cells) using a colorimetric assay that uses a reduction of a yellow tetrazolium salt MTT to purple formazan crystals by metabolically active cells. SH-SY5Y cells were maintained in Dulbecco’s modified Eagle’s medium/Nutrient Mixture F-12 Ham (DMEM F-12) while HepG2 was maintained in Dulbecco’s modified Eagle’s medium (DMEM), both supplemented with 10% fetal bovine serum (FBS). Monomeric (0 h incubation) and aggregated (192 h incubation) samples of NSP11 were re-suspended at the calculated volume of media to attain the desired concentration. Further, 6000 cells/well in a 96-well plate were seeded with 150 μl volume in culture media and incubated for 24 h at 37 °C to allow them to surface adhere. After incubation, cells were treated with varying concentrations of NSP11 (1.4 to 180 μM) samples in 100 μl fresh culture media and incubated at 37 °C for 24 h and 72 h. MTT (0.5 mg/ml final concentration) was added to each well and incubated for 3 h at 37 °C, and then 100 μl DMSO was added to dissolve formazan crystal (10 min incubation). The absorbance was recorded at 590 nm (630 nm reference wavelength) in TECAN Infinite M200 PRO multimode microplate reader. Data were collected in triplicates and reported as the relative percentage of cell viability with respect to untreated cells.

## Data availability

All data are contained within the manuscript or as supporting information.

## Acknowledgment

Authors are thankful to the Indian Institute of Technology Mandi (BioX, AMRC, and C4DFED center) for all the facilities and also to faculty research grant, School of Basic Sciences, IIT Mandi to RG. KG was supported by the Science and Engineering Research Board (SERB), India (Grant Number: CRG/2019/005603) for fellowship. KUS and SKK were supported by the Indian Council of Medical Research (ICMR), India, for fellowship. AK was supported by the Ministry of Human Resource Development (MHRD). TB is thankful to the Department of Science and Technology (DST) for the INSPire Fellowship. PK and NN were supported by CRG/2019/005603. RG is grateful for the IYBA Award (Grant Number: BT/11/IYBA/2018/06) from the Department of Biotechnology (DBT), India.

## Author contributions

RG, NG, and MV: Conception, design, review, data analysis and writing of the manuscript. TB, KG, PK, NN, SK, AK, ZFB and KUS performed experiments and computational analysis predictions. TB, KG, SKK, and PK analyzed data and wrote the manuscript.

## References

1. I. Chakraborty, P. Maity, COVID-19 outbreak: Migration, effects on society, global environment and prevention. Science of the Total Environment 728 (2020).

2. H. F. Florindo, et al., Immune-mediated approaches against COVID-19. Nature Nanotechnology 15, 630–645 (2020).

3. A. Moshe, R. Gorovits, Virus-Induced Aggregates in Infected Cells. Viruses 4, 2218–2232 (2012).

4. O. Olasunkanmi, S. Chen, J. Mageto, Z. Zhong, Virus-Induced Cytoplasmic Aggregates and Inclusions Are Critical Cellular Regulatory and Antiviral Factors. Viruses 12, 399 (2020).

5. C. Chevalier, et al., PB1-F2 influenza A virus protein adopts a β-sheet conformation and forms amyloid fibers in membrane environments. Journal of Biological Chemistry 285, 13233–13243 (2010).

6. C. L. Pham, et al., Viral M45 and necroptosis-associated proteins form heteromeric amyloid assemblies. EMBO reports 20 (2019).

7. K. Ezzat, et al., The viral protein corona directs viral pathogenesis and amyloid aggregation. Nature Communications 10 (2019).

8. A. Miodek, et al., Electrochemical detection of the oligomerization of PB1-F2 influenza a virus protein in infected cells. Analytical Chemistry 86, 9098–9105 (2014).

9. C. Chevalier, et al., Synchrotron infrared and deep UV fluorescent microspectroscopy study of PB1-F2 β-Aggregated structures in influenza a virus-infected cells. Journal of Biological Chemistry 291, 9060–9072 (2016).

10. S. Gao, et al., Characteristics of Nucleocytoplasmic Transport of H1N1 Influenza A Virus Nuclear Export Protein. Journal of Virology 88, 7455–7463 (2014).

11. A. O. Golovko, et al., Aggregation of Influenza A Virus Nuclear Export Protein. Biochemistry (Moscow) 83, 1411–1421 (2018).

12. A. Zlotnick, R. Aldrich, J. M. Johnson, P. Ceres, M. J. Young, Mechanism of capsid assembly for an icosahedral plant virus. Virology 277, 450–456 (2000).

13. Y. N. Lee, et al., Thermal aggregation of SARS-CoV membrane protein. Journal of Virological Methods 129, 152–161 (2005).

14. A. Ghosh, et al., Self-Assembly of a Nine-Residue Amyloid-Forming Peptide Fragment of SARS Corona Virus E-Protein: Mechanism of Self Aggregation and Amyloid-Inhibition of hIAPP. Biochemistry 54, 2249–2261 (2015).

15. J. L. Nieto-Torres, et al., Severe Acute Respiratory Syndrome Coronavirus Envelope Protein Ion Channel Activity Promotes Virus Fitness and Pathogenesis. PLoS Pathogens 10 (2014).

16. C. S. Shi, N. R. Nabar, N. N. Huang, J. H. Kehrl, SARS-Coronavirus Open Reading Frame-8b triggers intracellular stress pathways and activates NLRP3 inflammasomes. Cell Death Discovery 5 (2019).

17. R. Giri, et al., Understanding COVID-19 via comparative analysis of dark proteomes of SARS-CoV-2, human SARS and bat SARS-like coronaviruses. Cellular and Molecular Life Sciences (2020)https://doi.org/10.1007/s00018-020-03603-x.

18. L. Goldschmidt, P. K. Teng, R. Riek, D. Eisenberg, Identifying the amylome, proteins capable of forming amyloid-like fibrils. Proceedings of the National Academy of Sciences of the United States of America 107, 3487–3492 (2010).

19. A. K. Buell, C. M. Dobson, T. P. J. Knowles, M. E. Welland, Interactions between Amyloidophilic dyes and their relevance to studies of amyloid inhibitors. Biophysical Journal 99, 3492–3497 (2010).

20. P. Buzón, S. Maity, W. H. Roos, Physical virology: From virus self-assembly to particle mechanics. Wiley Interdisciplinary Reviews: Nanomedicine and Nanobiotechnology 12 (2020).

21. W. Kim, M. H. Hecht, Generic hydrophobic residues are sufficient to promote aggregation of the Alzheimer’s Aβ42 peptide. Proceedings of the National Academy of Sciences of the United States of America 103, 15824–15829 (2006).

22. A. L. Fink, Protein aggregation: Folding aggregates, inclusion bodies and amyloid. Folding and Design 3 (1998).

23. S. Satarker, M. Nampoothiri, Structural Proteins in Severe Acute Respiratory Syndrome Coronavirus-2. Archives of Medical Research 51, 482–491 (2020).

24. Y. Huang, C. Yang, X. feng Xu, W. Xu, S. wen Liu, Structural and functional properties of SARS-CoV-2 spike protein: potential antivirus drug development for COVID-19. Acta Pharmacologica Sinica 41, 1141–1149 (2020).

25. Z. F. Brotzakis, T. Lohr, M. Vendruscolo, Determination of Intermediate State Structures in the Opening Pathway of SARS-CoV-2 Spike Using Cryo-Electron Microscopy. ChemRxiv (2020) https://doi.org/10.26434/CHEMRXIV.13222073.V1.

26. M. Santerre, S. P. Arjona, C. N. Allen, N. Shcherbik, B. E. Sawaya, Why do SARS-CoV-2 NSPs rush to the ER? Journal of Neurology 1, 3 (2020).

27. M. I. Sulatsky, et al., Effect of the fluorescent probes ThT and ANS on the mature amyloid fibrils. Prion 14, 67–75 (2020).

28. C. Xue, T. Y. Lin, D. Chang, Z. Guo, Thioflavin T as an amyloid dye: Fibril quantification, optimal concentration and effect on aggregation. Royal Society Open Science 4 (2017).

29. K. L. Zapadka, F. J. Becher, A. L. Gomes dos Santos, S. E. Jackson, Factors affecting the physical stability (aggregation) of peptide therapeutics. Interface Focus 7 (2017).

30. J. D. Harper, P. T. Lansbury, Models of amyloid seeding in Alzheimer’s disease and scrapie: Mechanistic truths and physiological consequences of the time-dependent solubility of amyloid proteins. Annual Review of Biochemistry 66, 385–407 (1997).

31. X. Xia, Domains and Functions of Spike Protein in Sars-Cov-2 in the Context of Vaccine Design. Viruses 13, 109 (2021).

32. J. K. Millet, G. R. Whittaker, Physiological and molecular triggers for SARS-CoV membrane fusion and entry into host cells. Virology 517, 3–8 (2018).

33. K. Pancer, et al., The SARS-CoV-2 ORF10 is not essential in vitro or in vivo in humans. PLoS Pathogens 16, e1008959 (2020).

34. Y. Finkel, et al., The coding capacity of SARS-CoV-2. Nature 589, 125–130 (2020).

35. R. Cagliani, D. Forni, M. Clerici, M. Sironi, Coding potential and sequence conservation of SARS-CoV-2 and related animal viruses. Infection, Genetics and Evolution 83, 104353 (2020).

36. H. Xia, et al., Evasion of Type I Interferon by SARS-CoV-2. Cell Reports 33, 108234 (2020).

37. D. Benvenuto, et al., Evolutionary analysis of SARS-CoV-2: how mutation of Non-Structural Protein 6 (NSP6) could affect viral autophagy. Journal of Infection 81, e24–e27 (2020).

38. R. Giri, et al., Understanding COVID-19 via comparative analysis of dark proteomes of SARS-CoV-2, human SARS and bat SARS-like coronaviruses. Cellular and Molecular Life Sciences (2020) https://doi.org/10.1007/s00018-020-03603-x.

39. K. Gadhave, et al., NSP 11 of SARS-CoV-2 is an Intrinsically Disordered Protein. bioRxiv (2020) https://doi.org/10.1101/2020.10.07.330068.

40. A. Abbott, Are infections seeding some cases of Alzheimer’s disease? Nature 587, 22–25 (2020).

41. B. Niklasson, L. Lindquist, W. Klitz, E. Englund, Picornavirus Identified in Alzheimer’s Disease Brains: A Pathogenic Path? Journal of Alzheimer’s Disease Reports 4, 141–146 (2020).

42. D. B. Clifford, et al., CSF biomarkers of Alzheimer disease in HIV-associated neurologic disease. Neurology 73, 1982–1987 (2009).

43. K. Lopatko Lindman, et al., Herpesvirus infections, antiviral treatment, and the risk of dementia—a registry-based cohort study in Sweden. Alzheimer’s & Dementia: Translational Research & Clinical Interventions 7, e12119 (2021).

44. V. L. Nimgaonkar, et al., Temporal cognitive decline associated with exposure to infectious agents in a population-based, aging cohort. Alzheimer Disease and Associated Disorders 30, 216–222 (2016).

45. D. M. Cairns, et al., A 3D human brain–like tissue model of herpes-induced Alzheimer’s disease. Science Advances 6, eaay8828 (2020).

46. H. Jang, et al., Highly pathogenic H5N1 influenza virus can enter the central nervous system and induce neuroinflammation and neurodegeneration. Proceedings of the National Academy of Sciences of the United States of America 106, 14063–14068 (2009).

47. P. B. Mccray, et al., Lethal Infection of K18-hACE2 Mice Infected with Severe Acute Respiratory Syndrome Coronavirus. JOURNAL OF VIROLOGY 81, 813–821 (2007).

48. J. Xu, et al., Detection of severe acute respiratory syndrome coronavirus in the brain: Potential role of the chemokine Mig in pathogenesis. Clinical Infectious Diseases 41, 1089–1096 (2005).

49. L. K. Tsai, et al., Neuromuscular disorders in severe acute respiratory syndrome. Archives of Neurology 61, 1669–1673 (2004).

50. J. Netland, D. K. Meyerholz, S. Moore, M. Cassell, S. Perlman, Severe Acute Respiratory Syndrome Coronavirus Infection Causes Neuronal Death in the Absence of Encephalitis in Mice Transgenic for Human ACE2. JOURNAL OF VIROLOGY 82, 7264–7275 (2008).

51. N. Poyiadji, et al., COVID-19-associated acute hemorrhagic necrotizing encephalopathy: Imaging features. Radiology 296, E119–E120 (2020).

52. L. Mao, et al., Neurologic Manifestations of Hospitalized Patients with Coronavirus Disease 2019 in Wuhan, China. JAMA Neurology 77, 683–690 (2020).

53. L. Duong, P. Xu, A. Liu, “Meningoencephalitis without respiratory failure in a young female patient with COVID-19 infection in Downtown Los Angeles, early April 2020” (Academic Press Inc., 2020).

54. T. Moriguchi, et al., A first case of meningitis/encephalitis associated with SARS-Coronavirus-2. International Journal of Infectious Diseases 94, 55–58 (2020).

55. A. Ramani, et al., SARS-CoV-2targets neurons of 3D human brain organoids. The EMBO Journal 39, e106230 (2020).

56. O. Conchillo-Solé, et al., AGGRESCAN: A server for the prediction and evaluation of “hot spots” of aggregation in polypeptides. BMC Bioinformatics 8 (2007).

57. S. O. Garbuzynskiy, M. Y. Lobanov, O. V. Galzitskaya, FoldAmyloid: A method of prediction of amyloidogenic regions from protein sequence. Bioinformatics 26, 326–332 (2009).

58. M. Emily, A. Talvas, C. Delamarche, MetAmyl: A METa-predictor for AMYLoid proteins. PLoS ONE 8 (2013).

59. P. Gasior, M. Kotulska, FISH Amyloid - a new method for finding amyloidogenic segments in proteins based on site specific co-occurence of aminoacids. BMC Bioinformatics 15 (2014).

60. P. Sormanni, F. A. Aprile, M. Vendruscolo, The CamSol method of rational design of protein mutants with enhanced solubility. Journal of Molecular Biology 427, 478–490 (2015).

61. K. Gadhave, R. Giri, Amyloid formation by intrinsically disordered trans-activation domain of cMyb. Biochemical and Biophysical Research Communications 524, 446–452 (2020).

62. M. Biancalana, S. Koide, Molecular mechanism of Thioflavin-T binding to amyloid fibrils. Biochimica et Biophysica Acta - Proteins and Proteomics 1804, 1405–1412 (2010).

