## Supplementary Material for "Amyloidogenic proteins in the SARS-CoV and SARS-CoV-2 proteomes"

**Table S1:** Distribution of APRs in SARS-CoV-2 structural proteins.

| **SARS-CoV-2 protein (length in amino acids) (NCBI accession ID)** | **MetAmyl**  **(% of APRs)** | **FISH Amyloid**  **(% of APRs)** | **AGGRESCAN**  **(% of APRs)** | **FoldAmyloid**  **(% of APRs)** | **CamSol**  **(% of hydrophobicity)** |
| --- | --- | --- | --- | --- | --- |
| **Spike glycoprotein (1273)**  **(YP_009724390.1)** | 1-20, 28-33, 42-52, 58-73, 88-97, 105-110, 114-121, 124-132, 139-145, 166-176, 192-205, 226-236, 262-272, 284-289, 302-308, 311-336, 347-352, 357-384, 390-409, 428-437, 443-448, 472-477, 495-500, 507-517, 523-528, 534-544, 546-554, 583-600, 605-622, 635-645, 651-660, 670-675, 687-697, 703-708, 717-727, 729-744, 756-763, 767-772, 776-786, 801-806, 822-830, 855-861, 869-875, 879-885, 906-918, 939-944, 947-952, 958-963, 969-981, 991-998, 1003-1013, 1029-1037, 1056-1071, 1075-1081, 1092-1107, 1119-1137, 1159-1164, 1171-1181, 1221-1255, 1236-1273  **(51.76%)** | 35-39, 54-58, 89-93, 116-120, 125-129, 139-146, 167-171, 179-183, 200-204, 266-270, 325-329, 365-370, 373-377, 399-403, 406-410, 446-450, 469-473, 482-486, 502-506, 540-544, 550-554, 558-562, 591-599, 609-613, 689-693, 783-789, 822-826, 914-918, 953-957, 1001-1005, 1059-1068, 1127-1132, 1175-1180, 1206-1210, 1228-1232  **(15.47%)** | 1-8, 10-16, 31-38, 40-47, 50-56, 58-70, 86-91, 100-109, 114-133, 140-145, 162-167, 189-208, 231-247, 260-271, 303-311, 331-336, 338-354, 362-383, 390-400, 428-437, 448-457, 484-490, 508-520, 584-588, 590-599, 610-617, 666-671, 688-698, 716-727, 729-745, 751-759, 761-768, 783-787, 797-805, 817-833, 853-861, 864-870, 872-896, 898-914, 961-982, 1000-1013, 1044-1052, 1058-1068, 1097-1101, 1124-1137, 1171-1179, 1207-1212, 1214-1249, 1266-1273  **(45.01%)** | 1-10, 34-38, 53-69, 100-108, 116-120, 126-135, 141-146, 174-178, 190-195, 235-244, 264-269, 274-278, 327-331, 348-353, 366-370, 389-395, 431-436, 450-457, 487-492, 507-519, 539-543, 560-564, 582-586, 609-613, 691-697, 718-722, 738-743, 751-755, 780-784, 799-804, 819-825, 875-880, 894-907, 993-999, 1005-1014, 1047-1051, 1060-1067, 1101-1105, 1127-1132, 1210-1239  **(23.72%)** | 1-10, 55-68, 117-121, 125-134, 263-269, 431-436, 487-492, 508- 516, 540-544, 592-598, 608-613, 692-698, 717-724, 738-742, 752-761, 884-889, 895-899, 1004-1010, 1059- 1068, 1101-1106, 1127-1135, 1211-1242  **(15.16%)** |
| **Envelope (75)**  **(YP_009724392.1)** | 1-6, 9-63  **(81.3%)** | 10-14, 16-23, 25-29, 55-59  **(30.66%)** | 1-6, 11-53, 55-61  **(74.67%)** | 11-51, 56-60  **(61.34%)** | 12-34, 39-50, 54-59  **(54.6%)** |
| **Membrane (222)**  **(YP_009724393.1)** | 5-10, 21-40, 43-52, 60-82, 92-102, 118-123, 136-145, 167-173, 187-197, 208-213, 216-222  **(52.7%)** | 25-29, 44-56, 69-73, 99-103, 108-112, 201-205  **(17.11%)** | 19-38, 43-58, 60-108, 118-122, 124-131, 135-149, 171-181, 216-222  **(59%)** | 18-39, 44-60, 62-77, 81-105, 108-112, 118-122, 126-132, 138-147, 176-180, 197-202, 218-222  **(57.65%)** | 21-38, 46-79, 81-100  **(32.4%)** |
| **Nucleocapsid (419)**  **(YP_009724391.1)** | 12-17, 49-58, 108-113, 128-135, 154-162, 241-251, 265-275, 332-337, 346-355, 387-396, 412-417  **(22.19%)** | 70-75, 159-163, 238-244, 348-352, 385-389, 405-409  **(7.87%)** | 49-56, 107-116, 126-137, 156-161, 218-226, 310-322, 327-336, 348-356  **(18.37%)** | 51-55, 85-89, 106-113, 130-134, 156-160, 219-226, 313-317, 329-333, 350-355, 392-396  **(13.6%)** | 51-56, 107-113, 131-134, 156-160, 330-334  **(5.48%)** |

**Table S2:** Distribution of APRs in SARS-CoV-2 accessory proteins.

| **SARS-CoV-2 protein (length in amino acids) (NCBI accession ID)** | **MetAmyl**  **(% of APRs)** | **FISH Amyloid**  **(% of APRs)** | **AGGRESCAN**  **(% of APRs)** | **FoldAmyloid**  **(% of APRs)** | **CamSol**  **(% of APRs)** |
| --- | --- | --- | --- | --- | --- |
| **ORF3a (275)**  **(YP_009724397.2)** | 6-17, 31-36, 43-67, 72-97, 109-125, 153-158, 160- 172, 185-191, 196-207, 211-216, 220-237, 243-249, 251-260, 268-275,  **(62.99%)** | 44-53, 76-80, 86-90, 109-117, 119-125, 142-146, 166-170, 212-216, 224-233, 254-258  **(24%)** | 1-14, 31-35, 37-41, 43-67, 73-103, 105-132, 141-158, 160-171, 197-218, 227-236  **(80%)** | 1-10, 43-57, 77-97, 103-116, 118-133, 139-152, 154-159, 201-210, 212-217, 228-235, 244-248  **(45.45%)** | 5-13, 42-57, 78-98, 106-118, 127-131, 143-151, 165-169, 202-206, 213-217, 227-234  **(27.63%)** |
| **ORF3b (22)** | 8-22  **(68.18%)** | 9-17  **(40.9%)** | 4-22  **(86.36%)** | 1-22  **(100%)** | 2-22  **(95.4%)** |
| **ORF6 (61)**  **(YP_009724394.1)** | 4-41  **(62.29%)** | 7-11, 14-19, 28-37  **(34.4%)** | 1-39  **(63.93%)** | 1-10, 12-22, 24-38  **(59.01%)** | 1-21, 24-35  **(54.09%)** |
| **ORF7a (121)**  **(YP_009724395.1)** | 3-15, 23-32, 35-40, 55-66, 69-77, 93-115  **(60.33%)** | 5-9, 17-21, 100-111  **(18.18%)** | 1-18, 26-33, 55-66, 100-116  **(45.45%)** | 1-13, 16-22, 54-61, 73-77, 98-116  **(42.97%)** | 1-13, 56-66, 99-114  **(33.05%)** |
| **ORF7b (43)**  **(YP_009725296.1)** | 2-12, 18-31  **(58.1%)** | 12-28  **(39.53%)** | 1-32  **(74.41%)** | 1-32  **(74.41%)** | 7-32  **(60.46%)** |
| **ORF8 (121)**  **(YP_009724396.1)** | 1-17, 20-33, 37-42, 46-51, 57-67, 71-91, 95-105, 113-121  **(78.51%)** | 2-10, 76-80, 107-111  (**15.7%)** | 1-15, 39-49, 57-61, 71-84, 96-103, 114-121  **(50.41%)** | 1-12, 39-49, 58-62, 81-88, 97-104, 113-121  **(43.8%)** | 1-15, 78-88, 117-121  **(25.61%)** |
| **ORF9b (97)** | 18-24, 30-35, 37-47, 71-80, 89-97  **(44.32%)** | 40-45  (**6.18%)** | 20-24, 44-50, 74-79, 90-97  **(26.8%)** | 20-25, 42-48, 69-73, 91-95  **(23.71%)** | 71-76, 92-97  **(12.37%)** |
| **ORF10 (38)**  **(YP_009724396.1)** | 2-8, 11-18, 25-38  **(76.3%)** | 23-27, 31-35  (**23.68%)** | 1-9, 11-20, 25-38  **(86.84%)** | 1-20, 31-37  **(71.05%)** | 1-20, 32-38  **(63.15%)** |
| **ORF14 (73)** | 3-9, 27-42, 49-54, 65-73  **(52%)** | 39-44  **(8.21%)** | 34-39, 50-71  **(38.35%)** | 29-38, 51-72  **(43.83%)** | 2-7, 33-40, 59-72  **(38.35%)** |

**Table S3:** Distribution of APRs in SARS-CoV-2 non-structural proteins.

| **SARS-CoV-2 protein (length in amino acids) (NCBI accession ID)** | **MetAmyl**  **(% of APRs)** | **FISH Amyloid**  **(% of APRs)** | **AGGRESCAN**  **(% of APRs)** | **FoldAmyloid**  **(% of APRs)** | **CamSol**  **(% of APRs)** |
| --- | --- | --- | --- | --- | --- |
| **NSP1 (180)**  **(YP_009725297.1)** | 10-31, 50-61, 66-74, 84-89, 101-111, 114-119  **(36.66%)** | 59-66, 92-96, 105-114  **(12.77%)** | 1-5, 20-31, 52-58, 68-72, 84-93, 103-108, 110-114, 116-122  **(31.67%)** | 18-22, 25-29, 68-72, 83-88, 105-109, 116-124  **(19.44%)** | 15-20, 105-109  **(6.11%)** |
| **NSP2 (638)**  **(YP_009725298.1)** | 46-53, 57-62, 94-105, 118-123, 134-139, 155-161, 182-193, 223-238, 248-263, 278-288, 292-313, 318-334, 339-344, 362-372, 378-401, 415-431, 433-442, 468-489, 492-503, 509-519, 525-535, 575-581, 598-608  **(44%)** | 184-189, 257-261, 363-368, 424-428, 433-440, 458-462, 468-472, 494-498, 560-564  **(7.83%)** | 78-82, 97-105, 118-124, 136-142, 158-162, 182-190, 225-241, 282-286, 291-315, 353-371, 382-445, 447-451, 463-487, 494-520, 524-528, 557-561, 571-576, 603-614, 625-632  **(42.31%)** | 1-5, 48-52, 58-62, 92-96, 134-139, 186-191, 228-235, 241-245, 294-299, 362-370, 397-408, 415-421, 430-439, 460-464, 467-474, 496-501, 503-508, 603-612  **(19.43%)** | 92-98, 123-128, 186-192, 226-235, 295-299, 351-355, 414-431, 434-439, 482-486, 496-500, 504-508, 602- 608  **(12.85%)** |
| **NSP3 (1945)**  **(YP_009725299.1)** | 1-8, 10-17, 20-27, 39-48, 52-66, 102-108, 144-149, 166-175, 179-191, 198-212, 220-225, 234-242, 272-277, 279-291, 295-304, 331-336, 341-361, 367-374, 393-398, 417-429, 434-442, 451-462, 467-486, 504-512, 516-521, 526-545, 548-554, 570-585, 593-602, 604-620, 633-640, 652-670, 682-692, 702-707, 715-722, 730-737, 745-757, 759-768, 775-780, 783-789, 797-804, 808-813, 842-847, 863-868, 891-896, 899-908, 928-938, 942-950, 965-972, 975-981, 1002-1008, 1025-1031, 1045-1051, 1054-1060, 1062-1075, 1112-1117, 1136-1142, 1147-1155, 1171-1176, 1189-1194, 1206-1212, 1225-1230, 1243-1259, 1284-1289, 1319-1324, 1327-1343, 1346-1351, 1360-1367, 1380-1392, 1395-1400, 1404-1437, 1443-1448, 1453-1477, 1486-1499, 1501-1525, 1529-1543, 1549-1556, 1563-1595, 1603-1613, 1615-1623, 1634-1648, 1668-1687, 1699-1706, 1718-1728, 1738-1746, 1758-1773, 1775-1787, 1791-1798, 1810-1819, 1821-1826, 1828-1837, 1846-1852, 1857-1862, 1880-1885, 1928-1941  **(51.61%)** | 10-14, 20-24, 201-205, 209-213, 218-222, 371-375, 379-383, 434-439, 452-456, 476-480, 504-508, 533-537, 580-584, 596-600, 716-720, 746-750, 755-760, 785-789, 823-827, 946-950, 964-968, 1010-1014, 1068-1072, 1106-1110, 1148-1153, 1253-1258, 1335-1339, 1396-1400, 1409-1413, 1423-1428, 1433-1437, 1456-1460, 1509-1513, 1518-1522, 1537-1541, 1547-1551, 1571-1577, 1615-1619, 1644-1648, 1720-1724, 1776-1780, 1813-1817, 1855-1859  **(11.41%)** | 16-23, 42-46, 51-66, 68-73, 82-89, 98-106, 185-189, 201-219, 237-245, 248-252, 281-291, 295-301, 330-339, 341-365, 367-374, 417-421, 432-442, 453-458, 461-465, 468-483, 530-538, 550-558, 573-580, 582-586, 600-618, 632-637, 648-653, 667-672, 684-689, 735-740, 742-758, 763-771, 824-830, 843-848, 857-868, 891-903, 912-916, 928-935, 946-957, 986-991, 1029-1034, 1045-1050, 1062-1071, 1107-1112, 1137-1142, 1152-1157, 1172-1177, 1188-1199, 1252-1259, 1282-1286, 1298-1315, 1327-1351, 1353-1364, 1383-1398, 1404-1441, 1455-1468, 1470-1479, 1485-1557, 1559-1585, 1604-1608, 1610-1624, 1636-1647, 1671-1678, 1720-1727, 1738-1749, 1751-1758, 1770-1786, 1788-1793, 1795-1799, 1807-1820, 1833-1838, 1856-1861, 1888-1900, 1927-1945  **(42.5%)** | 83-90, 100-106, 244-248, 294-299, 324-329, 342-349, 353-359, 435-440, 479-483, 506-510, 534-540, 553-562, 602-608, 614-619, 632-637, 646-650, 694-698, 707-711, 748-755, 799-803, 814-818, 825-830, 856-866, 879-884, 891-899, 915-919, 930-935, 959-954, 1108-1115, 1139-1143, 1172-1176, 1189-1197, 1255-1259, 1276-1281, 1337-1345, 1351-1364, 1409-1428, 1433-1437, 1458-1463, 1488-1494, 1496-1505, 1507-1555, 1561-1582, 1591-1595, 1614-1619, 1630-1635, 1670-1674, 1683-1688, 1701-1705, 1720-1725, 1741-1756, 1813-1817, 1835-1839, 1854-1859, 1891-1896  **(22.51%)** | 21-25, 41-45, 53-59, 101-106, 199-205, 239-244, 534-540, 551-555, 604-608, 629-638, 752-756, 762-771, 858-865, 892-899, 915-920, 931-935, 1191-1195, 1348-1363, 1410-1429, 1432-1437, 1458-1467, 1470-1474, 1489-1505, 1507-1554, 1564-1583, 1720-1725, 1741-1746, 1777-1783, 1891-1895, 1933-1938  **(14.7%)** |
| **NSP4 (500)**  **(YP_009725300.1)** | 9-35, 39-44, 48-54, 57-62, 90-105, 109-114, 120-137, 147-152, 154-161, 166-171, 175-183, 208-218, 231-239, 253-266, 280-299, 311-320, 323-363, 373-389, 393-398, 402-412, 434-439, 444-449, 458-463, 483-488, 491-500  **(57.6%)** | 16-24, 40-44, 52-56, 117-121, 128-133, 257-261, 288-293, 297-301, 336-347, 358-362, 403-412, 419-423, 444-448, 484-491  **(18.2%)** | 1-28, 41-49, 90-103, 114-122, 124-135, 137-157, 170-181, 193-201, 203-217, 236-240, 253-269, 275-306, 309-332, 334-338, 340-370, 375-409, 411-427, 430-436, 438-453, 494-500  **(65.8%)** | 4-9, 11-29, 72-76, 89-97, 100-104, 118-127, 190-194, 211-215, 253-257, 262-271, 290-306, 312-333, 342-354, 358-371, 373-385, 389-396, 398-405, 417-422, 426-430, 434-440, 442-449  **(40%)** | 12-31, 90-94, 100-104, 119-124, 128-132, 253-258, 264-271, 286-302, 314-352, 360-385, 387-396, 417-421, 495-500  **(31.6%)** |
| **NSP5 (306)**  **(YP_009725301.1)** | 13-26, 33-47, 65-86, 89-94, 101-118, 123-128, 141-162, 166-176, 196-215, 223-228, 246-251, 254-262, 280-285, 296-306  **(56.2%)** | 103-107, 139-143, 155-159, 301-306  **(6.86%)** | 15-23, 30-38, 70-78, 80-90, 101-107, 109-121, 135-141, 143-166, 199-218, 221-225, 227-238, 254-269, 281-287, 295-301  **(51.3%)** | 29-44, 56-60, 74-78, 84-89, 102-106, 112-117, 154-165, 200-212, 217-222, 231-235, 260-264  **(27.45%)** | 16-24, 66-71, 112-117, 157-162, 199-212, 232-236, 302-306  **(16.6%)** |
| **NSP6 (290)**  **(YP_009725302.1)** | 15-31, 48-54, 72-82, 103-108, 113-129, 141-155, 165-226, 241-251, 275-290  **(55.8%)** | 26-30, 35-39, 41-45, 77-84, 90-94, 120-125, 150-154, 165-169, 171-181, 199-203, 211-219  **(23.79%)** | 10-43, 45-72, 74-86, 88-102, 109-129, 139-156, 161-197, 199-250, 269-281, 283-290  **(82.41%)** | 11-26, 31-45, 47-61, 64-73, 76-86, 88-100, 112-118, 120-127, 138-155, 162-170, 180-203, 208-246, 267-271, 273-277  **(67.24%)** | 12-39, 43-59, 66-72, 78-89, 91-95, 115-125, 141-155, 163-172, 176-185, 187-204, 207-232, 237-245  **(57.5%)** |
| **NSP7 (83)**  **(YP_009725303.1)** | 6-18, 28-33, 51-63  **(38.55%)** | 14-18  **(6%)** | 8-19, 29-33, 38-42, 50-63  **(43.37%)** | 10-16, 27-31, 37-41, 53-61  **(31.32%)** | 9-17, 30-34, 55-61  **(25.3%)** |
| **NSP8 (198)**  **(YP_009725304.1)** | 31-36, 44-49, 81-88, 102-107, 114-172, 182-190  **(42.92%)** | 129-133, 182-186  **(5%)** | 5-9, 11-16, 34-38, 86-96, 122-132, 148-158, 184-191  **(28.78%)** | 88-96, 115-122, 127-133, 151-156, 180-188  **(19.69%)** | 58-61, 89-93, 128-132, 147-151, 182-187  **(10.6%)** |
| **NSP9 (113)**  **(YP_009725305.1)** | 28-33, 61-68, 72-77, 85-90, 101-106  **(28.31%)** | - | 38-46, 64-68, 85-94, 100-110  **(30.97%)** | 29-33, 40-44, 51-55, 86-92  **(19.46%)** | 87-91  **(4.42%)** |
| **NSP10 (139)**  **(YP_009725306.1)** | 9-18, 337-47, 50-58, 72-81, 98-103, 107-123  **(45.32%)** | 15-19  **(3.59%)** | 10-22, 39-48, 54-58, 70-79, 94-99, 109-127  **(45.32%)** | 14-21, 42-47, 73-81, 117-122  **(20.86%)** | 13-20, 71-78, 116-122  **(15.1%)** |
| **NSP11 (13) (YP_009725312.1)** | 8-13  **(46.1%)** | 6-10  **(38.46%)** | 6-13  **(61.53%)** | - | 8-13  **(46.1%)** |
| **NSP12 (932)**  **(YP_009725308.1)** | 6-16, 25-32, 34-39, 41-47, 65-73, 75-81, 85-90, 111-116, 142-152, 171-176, 185-193, 198-207, 229-239, 243-249, 313-325, 329-347, 352-357, 359-369, 392-401, 404-411, 421-428, 435-440, 467-480, 490-498, 526-531, 535-540, 554-568, 577-592, 601-609, 632-638, 641-650, 659-664, 670-680, 689-710, 715-720, 762-790, 813-821, 826-831, 842-849, 855-865, 880-888, 925-932  **(44.2%)** | 6-10, 34-38, 77-81, 110-114, 146-150, 203-207, 329-333, 337-341, 352-356, 371-375, 491-495, 559-563, 569-573, 585-590, 671-675, 692-697, 775-779, 784-788, 828-832, 843-847, 884-888  **(11.48%)** | 9-19, 29-37, 39-53, 67-76, 123-132, 142-150, 173-177, 186-192, 199-207, 218-222, 233-242, 244-250, 265-271, 283-287, 310-321, 329-338, 340-351, 366-377, 396-406, 422-430, 436-440, 464-480, 530-535, 538-547, 557-566, 582-594, 596-606, 628-643, 647-652, 662-676, 689-705, 742-792, 838-847, 855-867, 879-888, 899-905, 919-926, 928-932  **(44.52%)** | 6-10, 30-37, 46-50, 67-73, 100-106, 120-133, 144-151, 172-176, 184-188, 190-194, 202-206, 214-218, 238-247, 266-275, 281-292, 306-319, 331-336, 346-350, 370-375, 394-400, 405-409, 438-443, 454-460, 467-474, 492-496, 504-508, 513-517, 526-530, 544-548, 628-639, 650-655, 664-668, 725-733, 743-751, 753-758, 763-768, 784-789, 791-795, 826-830, 842-846, 854-862, 879-890, 900-907, 921-926  **(33.15%)** | 201-205, 239-245, 308-323, 335-345, 395-399, 438-443, 469-474, 559-565, 691-701, 763-768, 784-791, 880-886, 902-906  **(10.4%)** |
| **NSP13 (601)**  **(YP_009725308.1)** | 1-11, 24-50, 55-65, 67-74, 79-101, 106-111, 120-126, 144-159, 179-185, 187-199, 205-214, 216-233, 235-240, 245-252, 262-267, 289-294, 300-310, 330-362, 366-376, 382-387, 394-401, 421-429, 447-457, 474-486, 491-501, 508-515, 520-526, 528-538, 541-551, 560-578, 587-592, 596-601  **(59%)** | 41-45, 191-195, 255-259, 266-270, 295-299, 395-399, 421-425, 478-482, 494-498, 542-546  **(8.31%)** | 1-11, 24-34, 36-46, 59-70, 83-95, 107-111, 133-137, 146-156, 177-186, 212-216, 219-233, 243-253, 290-299, 301-313, 319-323, 327-332, 351-363, 367-377, 380-384, 394-401, 420-433,448-454, 471-484, 495-501, 507-513, 522-525, 540-549, 565-578, 585-590  **(46.08%)** | 4-8, 18-33, 62-66, 69-73, 120-124, 163-167, 178-187, 223-228, 245-251, 291-300, 303-308, 353-359, 370-374, 395-399, 426-430, 453-457, 470-478, 497-501, 508-513, 542-547, 559-565, 571-577  **(24.45%)** | 2-9, 44-48, 65-69, 222-230, 252-256, 292-299, 305-309, 355-361, 543-547, 571-575  **(10.48%)** |
| **NSP14 (527)**  **(YP_009725309.1)** | 3-19, 24-33, 35-43, 100-105, 108-127, 159-171, 179-187, 194-202, 205-211, 214-223, 229-236, 240-245, 259-265, 275-280, 285-290, 311-321, 325-333, 346-351, 377-386, 394-402, 416-421, 432-441, 443-448, 455-479, 502-514  **(47%)** | 38-42, 114-118, 160-164, 194-201, 232-236, 380-384, 417-421, 459-463, 491-495  **(9.1%)** | 9-13, 15-19, 52-65, 82-92, 107-120, 152-157, 159-172, 178-202, 228-238, 240-246, 271-296, 298-303, 312-326, 363-372, 379-389, 400-404, 431-436, 438-442, 444-449, 461-466, 468-479, 495-515  **(45.73%)** | 50-60, 71-75, 81-89, 147-155, 159-169, 181-188, 196-200, 207-211, 225-229, 233-243, 245-249, 258-262, 274-282, 284-288, 290-294, 313-317, 348-352, 365-370, 381-387, 397-401, 403-407, 443-447, 491-497, 506-513, 515-522  **(33.20%)** | 114-122, 181-187, 224-230, 233-239, 381-387, 442-447, 503-512  **(10.24%)** |
| **NSP15 (346)**  **(YP_09725310.1)** | 7-14, 22-43, 51-56, 79-88, 94-106, 114-123, 129-134, 139-156, 162-197, 231-240, 273-280, 285-298, 301-306, 309-331, 336-342  **(56.93%)** | 23-28, 76-80, 82-86, 100-104, 119-123, 139-144, 164-169, 175-179, 184-188, 275-279, 301-307  **(17.34%)** | 1-7, 24-32, 35-39, 51-56, 68-77, 79-85, 96-106, 119-124, 143-147, 163-174, 180-184, 230-241, 245-256, 273-280, 289-304, 306-334  **(46.24%)** | 55-59, 83-88, 120-124, 132-136, 211-216, 222- 226, 230-238, 249-253, 277-281, 292-301, 326-333, 339-343  **(21.38%)** | 82-86, 98-104, 117-122, 248-252, 292-296, 319-323, 327-332  **(10.98%)** |
| **NSP16 (298)**  **(YP_009725311.1)** | 43-49, 53-58, 66-72, 78-84, 115-122, 124-131, 135-142, 148-158, 162-172, 190-198, 202-210, 225-230, 237-242, 258-263, 270-276, 286-295  **(42.28%)** | 27-31, 48-52, 109-113, 124-129, 204-208, 225-229, 278-282  **(12%)** | 38-61, 63-71, 84-89, 124-131, 149-161, 163-172, 179-197, 203-211, 222-233, 238-250, 252-261, 269-279, 290-294  **(50%)** | 15-23, 48-59, 63-70, 84-90, 124-130, 149-159, 185-195, 204-211, 221-232, 269-274  **(30.53%)** | 66-70, 150-156, 186-196, 206-210, 225-231  **(16.1%)** |

**Table S4:** Predicted APRs in structural proteins of SARS-CoV.

| **SARS-CoV Protein** **(length in amino acids) (UniProt ID)** | **MetAmyl**  **(% of APRs)** | **FISH Amyloid**  **(% of APRs)** | **AGGRESCAN**  **(% of APRs)** | **FoldAmyloid**  **(% of APRs)** | **CamSol**  **(Degree of hydrophobicity)** |
| --- | --- | --- | --- | --- | --- |
| **Spike Glycoprotein**  (1255)  (P59594) | 61-76, 87-92, 97-107, 111-130, 146-153, 159-173, 191-207, 211-216, 224-229, 231-239, 249-259, 271-276, 287-295, 298-311, 313-323, 328-339, 344-374, 377-394, 399-408, 419-424, 428-433, 456-461, 482-503, 509-514, 523-530, 532-540, 573-586, 594-612, 624-633, 637-646, 659-667, 672-679, 685-690, 700-706, 708-725, 738-745, 763-768, 783-788, 804-812, 837-843, 851-865, 888-900, 912-927, 929-934, 940-945, 951-963, 973-980, 985-995, 7011-1019, 1038-1053, 1057-1063, 1074-1087, 1101-1119, 1141-1146, 1153-1163, 1197-1227, 1230-1237, 1245-1255  **(52.19%)** | 3-10, 39-43, 58-62, 112-118, 160-164, 192-201, 252-256, 292-296, 352-357, 393-397, 403-407, 433-437, 500-504, 516-520, 523-530, 536-540, 545-549, 577-585, 595-599, 765-769, 804-808, 896-900, 935-939, 983-987, 1041-1050, 1075-1079, 1093-1097, 1109-1115, 1157-1162, 1188-1192, 1210-1214  **(14.42%)** | 1-12, 54-60, 62-68, 83-88, 97-106, 111-129, 137-142, 152-169, 189-201, 224-239, 247-258, 287-291, 310-316, 318-323, 325-334, 336-341, 349-370, 377-387, 415-425, 471-476, 478-492, 494-506, 508-512, 532-537, 576-585, 592-601, 637-641, 652-665, 669-685, 690-696, 698-709, 711-716, 718-723, 733-741, 743-747, 775-788, 799-815, 835-843, 846-852, 854-878, 880-896, 943-964, 982-995, 1026-1034, 1040-1050, 1073-1084, 1086-1093, 1095-1101, 1106-1119, 1153-1161, 1189-1194, 1196-1231, 1248-1255  **(53.46%)** | 1-10, 52-64, 99-103, 123-128, 150-155, 183-187, 193-198, 232-237, 251-256, 261-265, 314-318, 353-363, 376-381, 417-424, 438-444, 473-478, 493-503, 525-529, 595-599, 661-667, 675-679, 700-704, 720-725, 733-737, 762-766, 780-786, 801-807, 876-889, 975-981, 987-996, 1029-1033, 1042-1049, 1083-1087, 1090-1094, 1109-1114, 1192-1221  **(21.44%)** | 1-10, 59-66, 114-118, 158-162, 194-198, 250-256, 355-364, 418-422, 483-488, 494-501, 525-530, 578-584, 595-599, 657-663, 675-680, 697-706, 734-743, 854-858, 866-871, 877-881, 986-992, 1041-1050, 1076-1086, 1109-1114, 1193-1224  **(16.09%)** |
| **Envelope**  (76)  (P59637) | 1-6, 9-63  **(80.26%)** | 10-14, 16-23, 25-29  **(23.68%)** | 1-5, 11-53, 55-62  **(73.6%)** | 11-51, 56-60  **(60.53%)** | 12-37, 39-50, 54-59  **(57.89%)** |
| **Membrane**  (221)  (P59596) | 4-9, 20-27, 34-39, 42-53, 59-72, 74-81, 83-101, 117-132, 135-147, 166-172, 191-196, 215-221  **(55.2%)** | 24-28, 43-48, 68-72, 98-102, 107-111, 140-144, 200-204,  **(16.28%)** | 18-37, 42-57, 59-107, 117-121, 123-130, 134-147, 170-180, 216-221  **(58.37%)** | 17-38, 43-59, 61-75, 80-104, 107-111, 117-121, 137-146, 175-179, 196-201, 217-221  **(52.04%)** | 20-37, 45-78, 80-99  **(32.57%)** |
| **Nucleocapsid**  (422)  (P59595) | 13-18, 50-59, 109-114, 129-136, 156-162, 233-240, 242-252, 266-276, 333-338, 347-338, 347-356, 389-397  **(21.8%)** | 71-76, 160-164, 239-245, 349-353, 386-390  **(6.63%)** | 50-57, 108-117, 127-138, 218-227, 311-323, 328-336, 349-353  **(15.87%)** | 52-56, 86-90, 107-114, 131-135, 220-227, 314-318, 330-334, 351-356  **(11.14%)** | 52-57, 108-114, 131-135, 331-335  **(5.45%)** |

**Table S5:** Predicted APRs in accessory proteins of SARS-CoV.

| **SARS-CoV Proteins** **(length in amino acids) (UniProt ID)** | **MetAmyl**  **(% of APRs)** | **FISH Amyloid**  **(% of APRs)** | **AGGRESCAN**  **(% of APRs)** | **FoldAmyloid**  **(% of APRs)** | **CamSol**  **(% of APRs)** |
| --- | --- | --- | --- | --- | --- |
| **ORF3a**  (274)  (P59632) | 7-16, 27-36, 45-67, 76-97, 103-119, 143-152, 161-174, 185-190, 196-213, 216-221, 229-237, 242-248, 250-255, 267-274  **(60.58%)** | 49-55, 77-81, 86-91, 104-114, 142-146, 214-220, 228-233  **(17.15%)** | 1-16, 31-35, 37-41, 43-67, 69-132, 141-150, 160-168, 198-216, 227-236  **(59.48%)** | 1-10, 43-57, 61-65, 69-73, 75-97, 103-117, 119-132, 139-152, 154-159, 199-213, 229-235  **(47.08%)** | 4-8, 42-57, 76-98, 104-118, 126-131, 143-151, 199-211, 228-234  **(34.30%)** |
| **ORF3b**  (154)  (P59633) | 3-37, 43-61, 67-76, 79-87, 116-121, 137-148  **(59.09%)** | 16-25, 46-50, 116-121  **(13.63%)** | 4-34, 44-67, 71-88, 93-97, 116-123  **(55.84%)** | 4-8, 12-21, 25-31, 46-52, 61-65, 75-89, 112-125, 128-134  **(45.45%)** | 7-30, 46-51, 81-86, 116-122  **(27.92%)** |
| **ORF6**  (63)  (P59634) | 4-26, 28-41  **(58.73%)** | 7-11, 14-18, 29-34  **(25.39%)** | 1-39  **(61.90%)** | 1-10, 12-40  **(61.90%)** | 1-19, 24-35  **(49.20%)** |
| **ORF7a**  (122)  (P59635) | 3-17, 23-32, 56-64, 100-118  **(43.44%)** | 5-11, 17-21, 101-105, 108-113  **(18.85%)** | 1-18, 26-33, 55-67, 101-118  **(65.9%)** | 1-13, 16-22, 63-67, 73-77, 99-117  **(40.16%)** | 1-15, 56-66, 100-115  **(34.42%)** |
| **ORF7b**  (44)  (Q7TFA1) | 5-12, 18-35  **(59%)** | 12-28  **(38.63%)** | 5-33  **(65.9%)** | 5-32  **(63.64%)** | 7-32  **(59.09%)** |
| **ORF8a**  (39)  (Q7TFA0) | 1-25  **(61.53%)** | 2-6  **(12.8%)** | 1-21  **(53.8%)** | 1-22  **(54.41%)** | 2-21  **(51.28%)** |
| **ORF8b**  (84)  (Q80H93) | 2-10, 26-32, 34-49, 73-81  **(48.8%)** | 6-10  **(5.95%)** | 1-9, 17-28, 33-39, 54-63, 72-80  **(55.95%)** | 1-9, 18-23, 28-38, 56-64, 73-79  **(50%)** | 17-25, 74-78  **(16.66%)** |
| **ORF9b**  (98)  (P59636) | 4-9, 19-27, 41-48, 72-81, 90-98  **(42.8%)** | 41-46  **(6.12%)** | 12-17, 21-25, 45-50, 52-56, 75-80, 91-98  **(36.73%)** | 21-26, 43-49  **(13.27%)** | 73-79, 93-97  **(12.24%)** |
| **ORF14**  (70)  (Q7TLC7) | 35-44, 49-54, 58-66  **(35.71%)** | 36-44, 48-52  **(20%)** | 35-41, 44-48, 50-57, 59-64  **(37.14%)** | 35-39, 51-60  **(21.43%)** | 33-40  **(11.42%)** |

**Table S6:** Predicted APRs in non-structural proteins of SARS-CoV.

| **SARS-CoV (UniProt ID: P0C6X7) Proteins** **(length in amino acids)** | **MetAmyl**  **(% of APRs)** | **FISH Amyloid**  **(% of APRs)** | **AGGRESCAN**  **(% of APRs)** | **FoldAmyloid**  **(% of APRs)** | **CamSol**  **(% of APRs)** |
| --- | --- | --- | --- | --- | --- |
| **Nsp1**  (180) | 2-31, 50-55, 66-74, 82-90, 100-115  **(38.88%)** | 4-11, 59-66, 105-109  **(11.66%)** | 1-8, 20-31, 52-58, 68-74, 85-93, 98-108, 117-121  **(32.77%)** | 2-7, 18-22, 25-29, 68-72, 104-109,116-124  **(20%)** | 15-20, 103-109  **(7.22%)** |
| **Nsp2**  (638) | 1-6, 29-36, 46-53, 92-108, 118-123, 149-154, 166173, 182-190, 225-239, 279-288, 291-312, 318-332, 339-344, 348-354, 358-372, 384-394, 404-431, 437-442, 447-452, 454-462, 471-488, 494-506, 509-518, 525-533, 536-541, 557-564, 568-574, 576-581, 593-608  **(48.74%)** | 38-42, 100-104, 168-172, 292-297, 318-322, 326-330, 362-366, 413-418, 447-52, 458-462, 470-480, 511-515, 560-564,  **(11.59%)** | 97-104, 118-124, 137-141, 182-188, 226-242, 282-286, 291-324, 358-371, 384-393, 400-425, 433-445, 447-461, 464-484, 494-506, 509-515, 523-534, 572-576, 593-601, 603-612, 625-632  **(38.7%)** | 22-26, 48-52, 58-62, 92-96, 134-139, 150-158, 227-236, 241-245, 271-275, 285-289, 294-299, 365-370, 398-406, 415-420, 431-435, 445-452, 466-475, 496-501, 507-511, 527-531, 536-540, 548-552, 606-612, 631-635  **(23.2%)** | 227-236, 295-299, 413-424, 468-479, 496-501, 600-608  **(8.46%)** |
| **Nsp3**  (1922) | 1-10, 12-18, 21-28, 39-67, ,83-90, 103-109, 151-157, 185-191, 197-207, 213-219, 257-263, 273-279, 309-314, 320-337, 342-350, 372-384, 393-405, 444-450, 452-462, 482-488, 508-516, 525-530, 549-561, 569-584, 588-597, 603-616, 628-646, 628-646, 662-667, 681-686, 691-697, 707-712, 724-736, 738-743, 752-757, 760-766, 775-782, 8181-826, 833-840, 844-850, 868-885, 905-914, 918-927, 938-949, 952-967, 994-1001, 1018-1037, 1044-1051, 1102-1109, 1113-1119, 1124-1131, 1148-1162, 1182-1194, 1220-1237, 1240-1247, 1261-1266, 1279-1291, 1296-1302, 1306-1315, 1319-1345, 1359-1367, 1372-1378, 1383-1389, 1396-1416, 1420-1425, 1432-1438, 1444-1453, 1463-1473, 1475-1482, 1487-1492, 1501-1512, 1514-1519, 1525-1533, 1540-1560, 1562-1572, 1580-1590, 1592-1600, 1614-1625, 1645-1658, 1676-1683, 1695-1705, 1705-1723, 1735-1750, 1752-1765, 1769-1775, 1787-1814, 1823-1829, 1834-1839, 1857-1862, 1865-1871, 1897-1902, 1905-1917  **(49.58%)** | 5-9, 11-15, 196-205, 319-323, 331-335, 435-439, 455-459, 552-556, 577-581, 588-593, 682-686, 723-727, 762-766, 800-804, 932-927, 941-945, 987-991, 1045-1046, 1083-1087, 1115-1119, 1125-1130, 1149-1153, 1323-1327, 1373-1377, 1410-1414, 1548-1552, 1555-1559, 1592-1596, 1621-1625, 1697-1701, 1787-1791, 1799-1803, 1897-1871, 1909-1914,  **(9.26%)** | 43-47, 52-67, 83-91, 102-108, 144-148, 193-205, 215-222, 258-267, 273-279, 302-306, 308-317, 319-334, 345-351, 393-397, 408-418, 444-459, 480-485, 507-514, 526-534, 549-562, 571-587, 589-601, 608-613, 631-637, 643-647, 660-665, 695-699, 713-717, 719-734, 742-748, 774-780, 819-825, 833-846, 868-879, 905-912, 922-934, 937-944, 958-968, 1022-1032, 1039-1046, 1113-1119, 1129-1134, 1149-1154, 1165-1174, 1229-1235, 1258-1263, 1273-1292, 1294-1302, 1304-1310, 1323-1328, 1330-1341, 1360-1375, 1381-1418, 1440-1445, 1447-1456, 1462-1534, 1536-1562, 1564-1570, 1581-1585, 1587-1600, 1616-1624, 1648-1655, 1697-1704, 1715-1726, 1728-1735, 1747-1763, 1765-1776, 1783-1797, 1833-1838, 1866-1877, 1904-1922  **(41.4%)** | 23-27, 53-58, 84-92, 101-107, 200-205, 215-219, 289-293, 303-307, 323-328, 331-336, 343-350, 411-416, 443-447, 455-459, 510-514, 529-538, 547-559, 578-584, 591-595, 608-613, 622-626, 670-674, 683-687, 692-696, 728-732, 776-780, 791-796, 802-807, 833-843, 856-861, 868-876, 892-896, 907-912, 926-931, 1024-1028, 1116-1120, 1127-1133, 1149-1153, 1166-1174, 1232-1236, 1321-1326, 1328-1341, 1384-1407, 1409-1414, 1447-1452, 1477-1482, 1484-1501, 1504-1532, 1538-1559, 1568-1572, 1591-1596, 1609-1613, 1647-1651, 1660-1665, 1678-1682, 1697-1702, 1718-1733, 1790-1794, 1811-1816, 1831-1836, 1868-1873, 1897-1903  **(23.99%)** | 54-60, 85-89, 215-221, 322-328, 331-336, 510-514, 527-531, 577-583, 605-614, 744-748, 835-842, 869-875, 908-912, 943-947, 1150-1154, 1168-1172, 1326-1341, 1387-1414, 1447-1451, 1465-1472, 1476-1481, 1485-1503, 1505-1516, 1519-1531, 1541-1560, 1566-1570, 1697-1702, 1718-1723, 1831-1835  **(12.90%)** |
| **Nsp4**  (500) | 1-8, 13-36, 53-58, 86-104, 109-114, 120-137, 147-152, 154-161, 166-171, 208-218, 229-239, 250-278, 280-302, 307-312, 323-367, 372-387, 393-398, 402-412, 444-449, 483-488, 491-500  **(56.2%)** | 22-26, 52-56, 90-95, 117-121, 128-133, 229-233, 257-261, 264-268, 325-329, 336-347, 349-353, 358-362, 377-385, 406-410, 419-423, 444-448, 484-491  **(20.2%)** | 1-28, 30-35, 90-103, 113-122, 124-135, 137-159, 170-181, 193-201, 203-217, 230-240, 250-269, 275-307, 309-313, 315-332, 334-338, 340-370, 375-399, 401-427, 430-436, 439-453, 494-500  **(72.4%)** | 7-35, 72-76, 92-97, 99-104, 118-127, 190-194, 211-215, 253-257, 262-270, 290-307, 319-333, 342-354, 358-371, 373-387, 389-405, 417-422, 426-430, 436-440, 442-449  **(39.2%)** | 3-8, 14-33, 90-94, 119-124, 128-132, 253-258, 264-272, 286-302, 316-335, 337-354, 360-387, 390-395, 417-421, 495-500  **(31.4%)** |
| **Nsp5**  (306) | 13-26, 35-47, 63-82, 89-94, 101-118, 123-128, 141-162, 166-176, 196-215, 223-228, 246-251, 254-262, 280-287, 296-306  **(55.55%)** | 65-69, 103-107, 139-143, 155-159, 301-306  **(8.49%)** | 15-23, 31-35, 68-78, 80-84, 101-107, 109-121, 143-166, 199-218, 221-225, 227-238, 254-269, 281-287, 295-301  **(46.07%)** | 29-33, 36-44, 56-60, 74-78, 84-89, 102-106, 112-117, 154-165, 200-212, 217-222, 231-235, 260-264  **(26.8%)** | 16-24, 65-71, 112-117, 157-162, 201-212, 232-236, 302-306  **(16.33%)** |
| **Nsp6**  (290) | 6-11, 19-40, 45-52, 72-82, 113-118, 123-129, 141-155, 164-185, 187-197, 199-226, 241-251, 264-269, 283-290  **(55.51%)** | 17-30, 33-37, 41-45, 77-84, 90-94, 120-125, 150-154, 171-183, 211-219,  **(24.13%)** | 9-43, 45-72, 74-86, 88-102, 110-129, 139-156, 161-197, 199-250, 267-281, 283-290  **(83.44%)** | 11-26, 31-38, 40-45, 47-60, 64-73, 76-86, 88-99, 112-127, 138-155, 162-170, 180-203, 208-246, 267-271, 273-277  **(66.56%)** | 12-29, 31-40, 43-58, 66-73, 77-89, 91-95, 115-125, 141-153, 163-172, 176-204, 207-232, 237-245  **(57.93%)** |
| **Nsp7**  (83) | 6-18, 28-33, 51-63  **(38.55%)** | 14-18  **(6%)** | 8-19, 29-33, 38-42, 50-63  **(43.37%)** | 10-16, 27-31, 37-41, 53-61  **(31.33%)** | 9-17, 30-34, 55-61  **(25.30%)** |
| **Nsp8**  (198) | 31-36, 44-49, 81-88, 102-107, 114-124, 126-135, 145-172, 182-190  **(42.42%)** | 182-186  **(3.62%)** | 5-9, 11-15, 34-38, 86-96, 122-132, 148-158, 184-191  **(28.2%)** | 88-96, 115-122, 127-131, 151-156, 180-188  **(18.69%)** | 89-93, 128-133, 147-151, 182-187  **(11.11%)** |
| **Nsp9**  (113) | 28-33, 61-68, 72-77, 85-90, 101-106  **(28.31%)** | - | 39-45, 64-68, 85-94, 100-110  **(30.08%)** | 29-33, 40-44, 51-55, 86-92  **(19.47%)** | 87-91  **(4.42%)** |
| **Nsp10**  (139) | 9-18, 37-47, 50-58, 72-81, 98-103, 107-123  **(45.32%)** | 15-19  **(3.59%)** | 10-22, 39-48, 54-58, 70-79, 94-99, 113-127  **(42.44%)** | 14-20, 42-47, 73-81, 117-122  **(20.14%)** | 13-19, 73-78, 116-122  **(14.38%)** |
| **NSP11**  **(13)** | 4-13  **(76.9%)** | 6-10  **(38.46%)** | 6-13  **(61.53%)** | 6-10  **(38.46%)** | 8-13  **(46.1%)** |
| **Nsp12**  (932) | 4-16, 25-32, 34-39, 41-47, 66-73, ,85-91, 93-98, 111-116, 142-152, 171-176, 185-193, 198-207, 220-239, 243-49, 313-325, 329-347, 352-357, 359-369, 392-401, 404-411, 421-428, 435-440, 467-480, 490-498, 526-531, 535-540, 554-568, 577-592, 601-609, 632-638, 659-664, 670-680, 689-710, 715-720, 742-747, 761-769, 773-780, 782-790, 813-821, 842-849, 855-865, 880-888, 925-932  **(44.52%)** | 34-38, 77-81, 110-114, 146-150, 203-207, 230-234, 329-333, 337-341, 352-356, 371-375, 491-495, 559-563, 569-573, 585-590, 671-675, 692-697, 775-779, 784-788, 828-832, 843-847, 884-888  **(11.48%)** | 7-19, 29-37, 39-53, 66-75, 89-93, 97-103, 123-132, 142-150, 173-177, 187-192, 199-207, 218-226, 233-242, 244-250, 265-271, 282-287, 310-321, 329-338, 340-351, 366-377, 396-406, 422-430, 436-440, 464-480, 530-535, 538-547, 557-566, 582-594, 596-607, 628-638, 662-676, 689-705, 742-763, 773-792, 838-847, 855-867, 879-888, 899-905, 919-926, 928-932  **(44.7%)** | 6-10, 30-37, 46-50, 67-73, 99-105, 120-133, 144-151, 172-176, 190-194, 202-206, 214-218, 238-247, 266-275, 280-292, 306-319, 331-336, 346-350, 370-375, 394-400, 405-409, 438-443, 454-460, 467-474, 492-496, 504-508, 513-517, 526-530, 544-548, 628-639, 650-655, 664-668, 725-733, 743-751, 753-758, 783-789, 791-795, 826-830, 842-846, 854-862, 879-890, 900-907, 921-926  **(32.19%)** | 239-245, 308-320, 335-345, 395-399, 438-443, 469-474, 560-565, 691-701, 785-791, 880-886, 902-906  **(9.01%)** |
| **Nsp13**  (601) | 1-11, 24-50, 55-65, 67-74, 79-101, 106-111, 120-126, 144-159, 179-185, 187-199, 205-214, 216-233, 235-240, 245-252, 262-267, 289-294, 300-310, 330-336, 346-376, 382-387, 394-401, 421-429, 447-457, 474-486, 491-501, 508-515, 520-526, 528-538, 541-551, 560-568, 570-578, 587-592, 596-601  **(58.9%)** | 41-45, 191-195, 255-259, 266-270, 295-299, 395-399, 421-425, 478-482, 494-498, 542-546  **(8.31%)** | 1-11, 24-34, 36-46, 59-70, 83-95, 107-111, 133-137, 146-156, 177-186, 212-216, 219-233, 243-253, 290-299, 301-313, 319-323, 327-332, 351-363, 367-377, 380-384, 394-401, 420-433, 448-454, 471-484, 495-501, 507-513, 522-528, 540-549, 565-578, 585-590  **(46.08%)** | 4-8, 18-33, 62-66, 69-73, 120-124, 163-167, 178-187, 223-228, 245-251, 291-300, 303-308, 353-359, 370-374, 395-399, 426-430, 453-457, 470-478, 497-501, 508-513, 542-547, 559-565, 571-577  **(24.46%)** | 2-9, 44-48, 65-69, 222-230, 252-256, 292-299, 305-309, 354-361, 543-547, 571-575  **(10.48%)** |
| **Nsp14**  (527) | 3-19, 24-43, 100-105, 108-128, 159-171, 179-187, 194-202, 205-211, 214-224, 228-236, 240-245, 275-280, 285-290, 307-322, 325-333, 346-351, 377-386, 395-402, 416-421, 434-439, 443-448, 455-479, 502-512  **(56.9%)** | 38-42, 114-118, 160-164, 194-201, 232-236, 293-297, 380-384, 417-421, 459-463, 507-511  **(10.05%)** | 9-19, 28-38, 52-65, 82-92, 107-120, 152-157, 159-172, 178-202, 228-238, 240-246, 271-296, 312-326, 363-369, 379-389, 396-404, 444-449, 461-466, 468-479, 496-515  **(44.78%)** | 50-60, 71-75, 81-89, 147-155, 159-169, 181-188, 196-200, 225-229, 233-243, 245-249, 274-282, 284-288, 313-317, 348-352, 365-370, 381-387, 396-401, 403-407, 443-447, 472-477, 492-497, 506-513, 515-522  **(30.36%)** | 28-32, 114-122, 181-187, 226-230, 233-239, 380-387, 442-447, 503-512  **(10.81%)** |
| **Nsp15**  (346) | 5-14, 22-43, 51-56, 79-88, 94-106, 116-125, 129-134, 139-153, 160-187, 190-197, 231-240, 273-280, 285-298, 301-306, 309-331, 336-342  **(56.64%)** | 23-28, 34-43, 65-69, 76-80, 82-86, 95-104, 119-123, 139-144, 164-169, 175-179, 181-186, 275-279, 301-307  **(23.41%)** | 1-7, 24-33, 35-40, 51-56, 68-77, 79-85, 96-106, 116-124, 143-148, 162-177, 180-184, 216-222, 230-237, 245-254, 273-280, 289-304, 306-334  **(49.44%)** | 55-59, 83-88, 120-124, 132-136, 212-216, 222-226, 230-238, 249-253, 277-281, 292-301, 326-333, 339-343  **(21.09%)** | 82-86, 97-102, 118-122, 248-252, 291-296, 319-323  **(9.53%)** |
| **Nsp16**  (298) | 34-39, 43-49, 53-58, 66-72, 78-84, 115-122, 124-131, 148-157, 162-172, 190-198, 2020-208, 225-230, 237-242, 258-263, 270-275, 285-297  **(41.27%)** | 27-31, 48-52, 109-113, 124-129, 204-208, 225-229, 278-282,  **(9.04%)** | 38-61, 63-71, 84-89, 124-131, 149-161, 163-172, 179-197, 203-211, 225-233, 238-250, 252-260, 270-279, 290-294  **(48.32%)** | 15-23, 48-59, 63-70, 84-90, 124-130, 149-156, 185-195, 226-232, 269-274  **(25.17%)** | 50-62, 66-70, 150-156, 186-196, 225-231  **(14.42%)** |

**Table S7:** NetChop 3.1 prediction of 20S proteasome cleavage site in structural proteins of SARS-CoV-2. The red colored residues represent the cleavage site while the underlined and bold residues are APRs predicted by FISH Amyloid server.

| **Protein Name** | **Distribution of predicted 20s proteasome cleavage sites** | **Distribution of cleavage sites in aggregation prone regions (APR)** |
| --- | --- | --- |
| **Spike** | 1-MFVFLVLLPLVSSQCVNLTTRTQLPPAYTN-30  31-SFTR**GVYYP**DKVFRSSVLHSTQD**LFLPF**FS-60  61-NVTWFHAIHVSGTNGTKRFDNPVLPFND**GV**-90  91-**YFA**STEKSNIIRGWIFGTTLDSKTQ**SLLIV**-120  121-NNAT**NVVIK**VCEFQFCND**PFLGVYYH**KNNK-150  151-SWMESEFRVYSSANNC**TFEYV**SQPFLMD**LE**-180  181-**GKQ**GNFKNLREFVFKNIDG**YFKIY**SKHTPI-210  211-NLVRDLPQGFSALEPLVDLPIGINITRFQT-240  241-LLALHRSYLTPGDSSSGWTAGAAAY**YVGYL**-270  271-QPRTFLLKYNENGTITDAVDCALDPLSETK-300  301-CTLKSFTVEKGIYQTSNFRVQPTE**SIVRF**P-330  331-NITNLCPFGEVFNATRFASVYAWNRKRISN-360  361-CVAD**YSVLYN**SA**SFSTF**KCYGVSPTKLNDL-390  391-CFTNVYAD**SFVIR**GD**EVRQI**APGQTGKIAD-420  421-YNYKLPDDFTGCVIAWNSNNLDSKV**GGNYN**-450  451-YLYRLFRKSNLKPFERDI**STEIY**QAGSTPC-480  481-N**GVEGFN**CYFPLQSYGFQPTN**GVGYQ**PYRV-510  511-VVLSFELLHAPATVCGPKKSTNLVKNKCV**N**-540  541-**FNFN**GLTGT**GVLTE**SNK**KFLPF**QQFGRDIA-570  571-DTTDAVRDPQTLEILDIT**PCSFGGVSVIT**P-600  601-GTNTSNQV**AVLY**QDVNCTEVPVAIHADQLT-630  631-PTWRVYSTGSNVFQTRAGCLIGAEHVNNSY-660  661-ECDIPIGAGICASYQTQTNSPRRARSVA**SQ**-690  691-**SII**AYTMSLGAENSVAYSNNSIAIPTNFTI-720  721-SVTTEILPVSMTKTSVDCTMYICGDSTECS-750  751-NLLLQYGSFCTQLNRALTGIAVEQDKNTQE-780  781-VF**AQVKQIY**KTPPIKDFGGFNFSQILPDPS-810  811-KPSKRSFIEDL**LFNKV**TLADAGFIKQYGDC-840  841-LGDIAARDLICAQKFNGLTVLPPLLTDEMI-870  871-AQYTSALLAGTITSGWTFGAGAALQIPFAM-900  901-QMAYRFNGIGVTQ**NVLYE**NQKLIANQFNSA-930  931-IGKIQDSLSSTASALGKLQDVV**NQNAQ**ALN-960  961-TLVKQLSSNFGAISSVLNDILSRLDKVEAE-990 991-VQIDRLITGR**LQSLQ**TYVTQQLIRAAEIRAS-1020  1021-ANLAATKMSECVLGQSKRVDFCGKGYHLMS-1050  1051-FPQSAPHG**VVFLHVTYVP**AQEKNFTTAPAI-1080  1081-CHDGKAHFPREGVFVSNGTHWFVTQRNFYE-1110  1111-PQIITTDNTFVSGNCD**VVIGIV**NNTVYDPL-1140  1141-QPELDSFKEELDKYFKNHTSPDVDLGDISG-1170  1171-INAS**VVNIQK**EIDRLNEVAKNLNESLIDLQ-1200  1201-ELGKY**EQYIK**WPWYIWLGFIAGLIAIV**MVT**-1230  1231-**IM**LCCMTSCCSCLKGCCSCGSCCKFDEDDS-1260  1261-EPVLKGVKLHYT-1273 | Total sites: 196 (15%)  Sites within APR: 34 (17.34%)  Sites outside APR: 162 (82.65%) |
| **Envelope** | 1-MYSFVSEET**GTLIV**N**SVLLFLAF**V**VFLLV**T-30  31-LAILTALRLCAYCCNIVNVSLVKP**SFYVY**S-60  61-RVKNLNSSRVPDLLV-75 | Total sites: 17 (22%)  Sites within APR: 6 (35.30%)  Sites outside APR: 11 (64.70%) |
| **Membrane** | 1-MADSNGTITVEELKKLLEQWNLVI**GFLFL**T-30  31-WICLLQFAYANRN**RFLYIIKLIFLWL**LWPV-60  61-TLACFVLA**AVYRI**NWITGGIAIAMACLVGL-90  91-MWLSYFIA**SFRLF**ARTR**SMWSF**NPETNILL-120  121-**NVPL**HGTILTRPLLESELVIGAVILRGHLR-150  151-IAGHHLGRCDIKDLPKEITVATSRTLSYYK-180  181-LGASQRVAGDSGFAAYSRYRIGNYKLNTDH-210  211-SSSSDNIALLVQ-222 | Total sites: 58 (26%)  Sites within APR: 16 (27.59%)  Sites outside APR: 42 (72.41%) |
| **Nucleocapsid** | 1-MSDNGPQNQRNAPRITFGGPSDSTGSNQNG-30  31-ERSGARSKQRRPQGLPNNTASWFTALTQHG-60  61-KEDLKFPRG**QGVPIN**TNSSPDDQIGYYRRA-90  91-TRRIRGGDGKMKDLSPRWYFYYLGTGPEAG-120  121-LPYGANKDGIIWVATEGALNTPKDHIGTRN-150  151-PANNAAIV**LQLPQ**GTTLPKGFYAEGSRGGS-180  181-QASSRSSSRSRNSSRNSTPGSSRGTSPARM-210  211-AGNGGDAALALLLLDRLNQLESKMSGK**GQQ**-240  241-**QQGQ**TVTKKSAAEASKKPRQKRTATKAYNV-270  271-TQAFGRRGPEQTQGNFGDQELIRQGTDYKH-300  301-WPQIAQFAPSASAFFGMSRIGMEVTPSGTW-330  331-LTYTGAIKLDDKDPNFK**DQVIL**LNKHIDAY-360  361-KTFPPTEPKKDKKKKADETQALPQ**RQKKQ**Q-390  391-TVTLLPAADLDDFS**KQLQQ**SMSSADSTQA-419 | Total sites: 39 (9.30%)  Sites within APR: 4 (10.25%)  Sites outside APR: 35 (89.74%) |

**Table S8:** NetChop 3.1 prediction of 20S proteasome cleavage site in accessory proteins of SARS-CoV-2. The red colored residues represent the cleavage site while the underlined and bold residues are APRs predicted by FISH Amyloid server.

| **Protein Name** | **Distribution of predicted 20s proteasome cleavage sites** | **Distribution of cleavage sites in aggregation prone regions (APR)** |
| --- | --- | --- |
| **ORF3A** | 1-MDLFMRIFTIGTVTLKQGEIKDATPSDFVR-30  31-ATATIPIQASLPF**GWLIVGVALL**AVFQSAS-60  61-KIITLKKRWQLALSK**GVHFV**CNLLL**LFVTV**-90  91-YSHLLLVAAGLEAPFLYL**YALVYFLQS**I**NF**-120  121-**VRIIM**RLWLCWKCRSKNPLLY**DANYF**LCWH-150  151-TNCYDYCIPYNSVTS**SIVIT**SGDGTTSPIS-180  181-EHDYQIGGYTEKWESGVKDCVVLHSYFTSD-210  211-Y**YQLYS**TQLSTDT**GVEHVTFFIY**NKIVDEP-240  241-EEHVQIHTIDGSS**GVVNP**VMEPIYDEPTTT-270  271-TSVPL-275 | Total sites: 50 (18.18%)  Sites within APR: 17 (34.0%)  Sites outside APR: 33 (66.0%) |
| **ORF3B** | 1-MMPTIFFA**GILIVTTIV**YLTIV-22 | Total sites: 6 (27.2%)  Sites within APR: 2 (33.33%)  Sites outside APR: 4 (66.66%) |
| **ORF6** | 1-MFHLVD**FQVTI**AE**ILLIIM**RTFKVSIW**NLD**-30  31-**YIINLII**KNLSKSLTENKYSQLDEEQPMEI-60  61-D | Total sites: 10 (16.39%)  Sites within APR: 6 (60.0%)  Sites outside APR: 4 (40%) |
| **ORF7A** | 1-MKII**LFLAL**ITLATCE**LYHYQ**ECVRGTTVL-30  31-LKEPCSSGTYEGNSPFHPLADNKFALTCFS-60  61-TQFAFACPDGVKHVYQLRARSVSPKLFIRQ-90  91-EEVQELYSP**IFLIVAAIVFIT**LCFTLKRKT-120  121-E | Total sites: 20 (16.52%)  Sites within APR: 9 (45.0%)  Sites outside APR: 11 (55.0%) |
| **ORF7B** | 1-MIELSLIDFYL**CFLAFLLFLVLIMLIIF**WF-30  31-SLELQDHNETCHA-43 | Total sites: 19 (44.18%)  Sites within APR: 12 (63.16%)  Sites outside APR: 7 (36.84%) |
| **ORF8** | 1-M**KFLVFLGII**TTVAAFHQECSLQSCTQHQP-30  31-YVVDDPCPIHFYSKWYIRVGARKSAPLIEL-60  61-CVDEAGSKSPIQYIDI**GNYTV**SCLPFTINC-90  91-QEPKLGSLVVRCSFYE**DFLEY**HDVRVVLDF-120  121-I | Total sites: 19 (15.70%)  Sites within APR: 5 (26.31%)  Sites outside APR:14 (73.68%) |
| **ORF9B** | 1-MDPKISEMHPALRLVDPQIQLAVTRMENAV-30  31-GRDQNNVGP**KVYPII**LRLGSPLSLNMARKT-60  61-LNSLEDKAFQLTPIAVQMTKLATTEELPDE-90  91-FVVVTVK-97 | Total sites: 12 (12.37%)  Sites within APR: 1 (8.33%)  Sites outside APR:11 (91.66%) |
| **ORF10** | 1-MGYINVFAFPFTIYSLLLCRMN**SRNYI**AQV-30  31-**DVVN**FNLT-38 | Total sites: 9 (23.68%)  Sites within APR: 2 (22.22%)  Sites outside APR:7 (77.77%) |
| **ORF14** | 1-MLQSCYNFLKEQHCQKASTQKGAEAAVKPL-30  31-LVPHHVVA**TVQEIQ**LQAAVGELLLLEWLAM-60  61-AVMLLLLCCCLTD-73 | Total sites: 13 (17.80%)  Sites within APR: 1 (7.69%)  Sites outside APR: 12 (92.31%) |

**Table S9:** NetChop 3.1 prediction of 20S proteasome cleavage site in non-structural proteins of SARS-CoV-2. The red colored residues represent the cleavage site while the underlined and bold residues are APRs predicted by FISH Amyloid server.

| **Protein Name** | **Distribution of predicted 20s proteasome cleavage sites** | **Distribution of cleavage sites in aggregation prone regions (APR)** |
| --- | --- | --- |
| **NSP1** | 1-MESLVPGFNEKTHVQLSLPVLQVRDVLVRG-30  31-FGDSVEEVLSEARQHLKDGTCGLVEVEK**GV**-60  61-**LPQLEQ**PYVFIKRSDARTAPHGHVMVELVA-90  91-E**LEGIQ**YGRSGET**LGVLVPHVGEI**PVAYRK-120  121-VLLRKNGNKGAGGHSYGADLKSFDLGDELG-150  151-TDPYEDFQENWNTKHSSGVTRELMRELNGG-180 | Total sites: 19 (10.5%)  Sites within APR: 1 (5.26%)  Sites outside APR: 18 (94.74%) |
| **NSP2** | 1-AYTRYVDNNFCGPDGYPLECIKDLLARAGK-30  31-ASCTLSEQLDFIDTKRGVYCCREHEHEIAW-60  61-YTERSEKSYELQTPFEIKLAKKFDTFNGEC-90  91-PNFVFPLNSIIKTIQPRVEKKKLDGFMGRI-120  121-RSVYPVASPNECNQMCLSTLMKCDHCGETS-150  151-WQTGDFVKATCEFCGTENLTKEGATTCGYL-180  181-PQN**AVVKIY**CPACHNSEVGPEHSLAEYHNE-210  211-SGLKTILRKGGRTIAFGGCVFSYVGCHNKC-240  241-AYWVPRASANIGCNHT**GVVGE**GSEGLNDNL-270  271-LEILQKEKVNINIVGDFKLNEEIAIILASF-300  301-SASTSAFVETVKGLDYKAFKQIVESCGNFK-330  331-VTKGKAKKGAWNIGEQKSILSPLYAFASEA-360  361-AR**VVRSIF**SRTLETAQNSVRVLQKAAITIL-390  391-DGISQYSLRLIDAMMFTSDLATNNLVVMAY-420  421-ITG**GVVQL**TSQW**LTNIFGTV**YEKLKPVLDW-450  451-LEEKFKE**GVEFL**RDGWE**IVKFI**STCACEIV-480  481-GGQIVTCAKEIKE**SVQTF**FKLVNKFLALCA-510  511-DSIIIGGAKLKALNLGETFVTHSKGLYRKC-540  541-VKSREETGLLMPLKAPKEI**IFLEG**ETLPTE-570  571-VLTEEVVLKTGDLQPLEQPTSEAVEAPLVG-600  601-TPVCINGLMLLEIKDTEKYCALAPNMMVTN-630  631-NTFTLKGG-638 | Total sites: 76 (11.91%)  Sites within APR: 7 (9.21%)  Sites outside APR:69 (90.79%) |
| **NSP3** | 1-APTKVTFGD**DTVIE**VQGYK**SVNIT**FELDER-30  31-IDKVLNEKCSAYTVELGTEVNEFACVVADA-60  61-VIKTLQPVSELLTPLGIDLDEWSMATYYLF-90  91-DESGEFKLASHMYCSFYPPDEDEEEGDCEE-120  121-EEFEPSTQYEYGTEDDYQGKPLEFGATSAA-150  151-LQPEEEQEEDWLDDDSQQTVGQQDGSEDNQ-180  181-TTTIQTIVEVQPQLEMELTP**VVQTI**EVN**SF**-210  211-**SGY**LKLT**DNVYI**KNADIVEEAKKVKPTVVV-240  241-NAANVYLKHGGGVAGALNKATNNAMQVESD-270  271-DYIATNGPLKVGGSCVLSGHNLAKHCLHVV-300  301-GPNVNKGEDIQLLKSAYENFNQHEVLLAPL-330  331-LSAGIFGADPIHSLRVCVDTVRTNVYLAVF-360  361-DKNLYDKLVS**SFLEM**KSE**KQVEQ**KIAEIPK-390  391-EEVKPFITESKPSVEQRKQDDKKIKACVEE-420  421-VTTTLEETKFLTE**NLLLYI**DINGNLHPDSA-450  451-T**LVSDI**DITFLKKDAPYIVGDVVQE**GVLTA**-480  481-VVIPTKKAGGTTEMLAKALRKVP**TDNYI**TT-510  511-YPGQGLNGYTVEEAKTVLKKCK**SAFYI**LPS-540  541-IISNEKQEILGTVSWNLREMLAHAEETRKL-570  571-MPVCVETKA**IVSTI**QRKYKGIKIQE**GVVDY**-600  601-GARFYFYTSKTTVASLINTLNDLNETLVTM-630  631-PLGYVTHGLNLEEAARYMRSLKVPATVSVS-660  661-SPDAVTAYNGYLTSSSKTPEEHFIETISLA-690  691-GSYKDWSYSGQSTQLGIEFLKRGDK**SVYYT**-720  721-SNPTTFHLDGEVITFDNLKTLLSLR**EVRTI**-750  751-KVFT**TVDNI**NLHTQVVDMSMTYGQQFGPTY-780  781-LD**GADVT**KIKPHNSHEGKTFYVLPNDDTLR-810  811-VEAFEYYHTTDPSFLGRYMSALNHTKKWKY-840  841-PQVNGLTSIKWADNNCYLATALLTLQQIEL-870  871-KFNPPALQDAYYRARAGEAANFCALILAYC-900  901-NKTVGELGDVRETMSYLFQHANLDSCKRVL-930  931-NVVCKTCGQQQTTLK**GVEAV**MYMGTLSYEQ-960  961-FKK**GVQIP**CTCGKQATKYLVQQESPFVMMS-990  991-APPAQYELKHGTFTCASEYT**GNYQC**GHYKH-1020  1021-ITSKETLYCIDGALLTKSSEYKGPITDVFY-1050  1051-KENSYTTTIKPVTYKLD**GVVCT**EIDPKLDN-1080  1081-YYKKDNSYFTEQPIDLVPNQPYPNA**SFDNF**-1110  1111-KFVCDNIKFADDLNQLTGYKKPASRELKVT-1140  1141-FFPDLNG**DVVAID**YKHYTPSFKKGAKLLHK-1170  1171-PIVWHVNNATNKATYKPNTWCIRCLWSTKP-1200  1201-VETSNSFDVLKSEDAQGMDNLACEDLKPVS-1230  1231-EEVVENPTIQKDVLECNVKTT**EVVGDI**IL-1260  1261-KPANNSLKITEEVGHTDLMAAYVDNSSLTI-1290  1291-KKPNELSRVLGLKTLATHGLAAVNSVPWDT-1320  1321-IANYAKPFLNKVVS**TTTNI**VTRCLNRVCTN-1350  1351-YMPYFFTLLLQLCTFTRSTNSRIKASMPTT-1380  1381-IAKNTVKSVGKFCLE**ASFNY**LKSPNFSK**LI**-1410  1411-**NII**IWFLLLSVC**LGSLIY**STAA**LGVLM**SNL-1440  1441-GMPSYCTGYREGYLN**STNVT**IATYCTGSIP-1470  1471-CSVCLSGLDSLDTYPSLETIQITISSFKWD1500  1501-LTAFGLVAE**WFLAY**ILFT**RFFYV**LGLAAIM-1530  1531-QLFFSY**FAVHF**ISNSW**LMWLI**INLVQMAPI-1560  1561-SAMVRMYIFF**ASFYYVW**KSYVHVVDGCNSS-1590  1591-TCMMCYKRNRATRVECTTIVNGVR**RSFYV**Y-1620  1621-ANGGKGFCKLHNWNCVNCDTFCA**GSTFI**SD-1650  1651-EVARDLSLQFKRPINPTDQSSYIVDSVTVK-1680  1681-NGSIHLYFDKAGQKTYERHSLSHFVNLDNL-1710  1711-RANNTKGSL**PINVI**VFDGKSKCEESSAKSA-1740  1741-SVYYSQLMCQPILLLDQALVSDVGDSAEVA-1770  1771-VKMFD**AYVNT**FSSTFNVPMEKLKTLVATAE-1800  1801-AELAKNVSLDNV**LSTFI**SAARQGFVDSDVE-1830  1831-TKDVVECLKLSHQSDIEVTGDSCN**NYMLT**Y-1860  1861-NKVENMTPRDLGACIDCSARHINAQVAKSH-1890  1891-NIALIWNVKDFMSLSEQLRKQIRSAAKKNN-1920  1921-LPFKLTCATTRQVVNVVTTKIALKGG-1946 | Total sites: 293 (15.05%)  Sites within APR: 51 (17.40%)  Sites outside APR: 242 (82.60%) |
| **NSP4** | 1-KIVNNWLKQLIKVTL**VFLFVAAIF**YLITPV-30  31-HVMSKHTDF**SSEII**GYKAIDG**GVTRD**IAST-60  61-DTCFANKHADFDTWFSQRGGSYTNDKACPL-90  91-IAAVITREVGFVVPGLPGTILRTTNG**DFLH**-120  121-**F**LPRVFS**AVGNIC**YTPSKLIEYTDFATSAC-150  151-VLAAECTIFKDASGKPVPYCYDTNVLEGSV-180  181-AYESLRPDTRYVLMDGSIIQFPNTYLEGSV-210  211-RVVTTFDSEYCRHGTCERSEAGVCVSTSGR-240  241-WVLNNDYYRSLPGVFC**GVDAV**NLLTNMFTP-270  271-LIQPIGALDISASIVAG**GIVAIV**VTC**LAYY**-300  301-**F**MRFRRAFGEYSHVVAFNTLLFLMSFTVLC-330  331-LTPVY**SFLPGVYSVIYL**YLTFYLTNDV**SFL**-360  361-**AH**IQWMVMFTPLVPFWITIAYIICISTKHF-390  391-YWFFSNYLKRRV**VFNGVSFSTF**EEAALC**TF**-420  421-**LLN**KEMYLKLRSDVLLPLTQYNR**YLALY**NK-450  451-YKYFSGAMDTTSYREAACCHLAKALNDFSN-480  481-SGS**DVLYQPPQ**TSITSAVLQ-500 | Total sites: 109 (21.8%)  Sites within APR: 20 (18.34%)  Sites outside APR: 89 (81.66%) |
| **NSP5** | 1-SGFRKMAFPSGKVEGCMVQVTCGTTTLNGL-30  31-WLDDVVYCPRHVICTSEDMLNPNYEDLLIR-60  61-KSNHNFLVQAGNVQLRVIGHSMQNCVLKLK-90  91-VDTANPKTPKYK**FVRIQ**PGQTFSVLACYNG-120  121-SPSGVYQCAMRPNFTIKG**SFLNG**SCGSVGF-150  151-NIDY**DCVSF**CYMHHMELPTGVHAGTDLEGN-180  181-FYGPFVDRQTAQAAGTDTTITVNVLAWLYA-210  211-AVINGDRWFLNRFTTTLNDFNLVAMKYNYE-240  241-PLTQDHVDILGPLSAQTGIAVLDMCASLKE-270  271-LLQNGMNGRTILGSALLEDEFTPFDVVRQC-300  301-**SGVTFQ**-306 | Total sites: 45 (14.70%)  Sites within APR: 4 (8.88%)  Sites outside APR: 41 (91.12%) |
| **NSP6** | 1-SAVKRTIKGTHHWLLLTILTSLLVL**VQSTQ**-30  31-WSLF**FFLYE**N**AFLPF**AMGIIAMSAFAMMFV-60  61-KHKHAFLCLFLLPSLA**TVAYFNMV**YMPAS**W**-90  91-**VMRI**MTWLDMVDTSLSGFKLKDCVMYASA**V**-120  121-**VLLIL**MTARTVYDDGARRVWTLMNVLTLV**Y**-150  151-**KVYY**GNALDQAISM**WALII**S**VTSNYSGVVT**-180  181-**T**VMFLARGIVFMCVEYCP**IFFIT**GNTLQCI-210  211-**MLVYCFLGY**FCTCYFGLFCLLNRYFRLTLG-240  241-VYDYLVSTQEFRYMNSQGLLPPKNSIDAFK-270  271-LNIKLLGVGGKPCIKVATVQ-290 | Total sites: 92 (31.72%)  Sites within APR: 30 (32.60%)  Sites outside APR: 62 (67.40%) |
| **NSP7** | 1-SKMSDVKCTSVVL**LSVLQ**QLRVESSSKLWA-30  31-QCVQLHNDILLAKDTTEAFEKMVSLLSVLL-60  61-SMQGAVDINKLCEEMLDNRATLQ-83 | Total sites: 9 (10.84%)  Sites within APR: 1 (11.11%)  Sites outside APR: 8 (88.89%) |
| **NSP8** | 1-AIASEFSSLPSYAAFATAQEAYEQAVANGD-30  31-SEVVLKKLKKSLNVAKSEFDRDAAMQRKLE-60  61-KMADQAMTQMYKQARSEDKRAKVTSAMQTM-90  91-LFTMLRKLDNDALNNIINNARDGCVPLNII-120  121-PLTTAAKL**MVVIP**DYNTYKNTCDGTTFTYA-150  151-SALWEIQQVVDADSKIVQLSEISMDNSPNL-180  181-A**WPLIV**TALRANSAVKLQ-198 | Total sites: 20 (10.10%)  Sites within APR: 2 (10.0%)  Sites outside APR: 18 (90.0%) |
| **NSP9** | 1-NNELSPVALRQMSCAAGTTQTACTDDNALA-30  31-YYNTTKGGRFVLALLSDLQDLKWARFPKSD-60  61-GTGTIYTELEPPCRFVTDTPKGPKVKYLYF-90  91-IKGLNNLNRGMVLGSLAATVRLQ-113 | Total sites: 16 (14.15%)  Sites within APR: 0 (0.0%)  Sites outside APR: 16 (100%) |
| **NSP10** | 1-AGNATEVPANSTVL**SFCAF**AVDAAKAYKDY-30  31-LASGGQPITNCVKMLCTHTGTGQAITVTPE-60  61-ANMDQESFGGASCCLYCRCHIDHPNPKGFC-90  91-DLKGKYVQIPTTCANDPVGFTLKNTVCTVC-120  121-GMWKGYGCSCDQLREPMLQ-139 | Total sites: 17 (12.23%)  Sites within APR: 2 (11.76%)  Sites outside APR: 15 (88.24%) |
| **NSP11** | 1-SADAQ**SFLNG**FAV-13 | Total sites: 2 (15.38%)  Sites within APR: 1 (50%)  Sites outside APR: 1 (50%) |
| **NSP12** | 1-SADAQ**SFLNR**VCGVSAARLTPCGTGTSTDV-30  31-VYR**AFDIY**NDKVAGFAKFLKTNCCRFQEKD-60  61-EDDNLIDSYF**VVKRHTFS**NYQHEETIYNLL-90  91-KDCPAVAKHDFFKFRIDGD**MVPHI**SRQRLT-120  121-KYTMADLVYALRHFDEGNCDTLKEI**LVTYN**-150  151-CCDDDYFNKKDWYDFVENPDILRVYANLGE-180  181-RVRQALLKTVQFCDAMRNAGIV**GVLTL**DNQ-210  211-DLNGNWYDFGDFIQTTPGSGVPVVDSYYSL-240  241-LMPILTLTRALTAESHVDTDLTKPYIKWDL-270  271-LKYDFTEERLKLFDRYFKYWDQTYHPNCVN-300  301-CLDDRCILHCANFNVLFSTVFPPTSFGP**LV**-330  331-**RKI**FVD**GVPFV**VSTGYHFREL**GVVHN**QDVN-360  361-LHSSRLSFKE**LLVYA**ADPAMHAASGNLLLD-390  391-KRTTCFSVAALTNNVAFQTVKPGNFNKDFY-420  421-DFAVSKGFFKEGSSVELKHFFFAQDGNAAI-450  451-SDYDYYRYNLPTMCDIRQLLFVVEVVDKYF-480  481-DCYDGGCINA**NQVIV**NNLDKSAGFPFNKWG-510  511-KARLYYDSMSYEDQDALFAYTKRNVIPTIT-540  541-QMNLKYAISAKNRARTVA**GVSIC**STMTN**RQ**-570  571-**FHQ**KLLKSIAATRG**ATVVIG**TSKFYGGWHN-600  601-MLKTVYSDVENPHLMGWDYPKCDRAMPNML-630  631-RIMASLVLARKHTTCCSLSHRFYRLANECA-660  661-QVLSEMVMCG**GSLYV**KPGGTSSGDATTAYA-690  691-N**SVFNIC**QAVTANVNALLSTDGNKIADKYV-720  721-RNLQHRLYECLYRNRDVDTDFVNEFYAYLR-750  751-KHFSMMILSDDAVVCFNSTYASQG**LVASI**K-780  781-NFK**SVLYY**QNNVFMSEAKCWTETDLTKGPH-810  811-EFCSQHTMLVKQGDDYV**YLPYP**DPSRILGA-840  841-GC**FVDDI**VKTDGTLMIERFVSLAIDAYPLT-870  871-KHPNQEYADVFHLYLQYIRKLHD**ELTGH**ML-900  901-DMYSVMLTNDNTSRYWEPEFYEAMYTPHTV-930  931-LQ-932 | Total sites: 155 (16.63%)  Sites within APR: 21 (13.54%)  Sites outside APR: 134 (86.46%) |
| **NSP13** | 1-AVGACVLCNSQTSLRCGACIRRPFLCCKCC-30  31-YDHVISTSHK**LVLSV**NPYVCNAPGCDVTDV-60  61-TQLYLGGMSYYCKSHKPPISFPLCANGQVF-90  91-GLYKNTCVGSDNVTDFNAIATCDWTNAGDY-120  121-ILANTCTERLKLFAAETLKATEETFKLSYG-150  151-IATVREVLSDRELHLSWEVGKPRPPLNRNY-180  181-VFTGYRVTKNSKVQIGEYTF**EKGDY**GDAVV-210  211-YRGTTTYKLNVGDYFVLTSHTVMPLSAPTL-240  241-VPQEHYVRITGLYP**TLNIS**DEFSSN**VANYQ**-270  271-KVGMQKYSTLQGPPGTGKSHFAIG**LALYY**P-300  301-SARIVYTACSHAAVDALCEKALKYLPIDKC-330  331-SRIIPARARVECFDKFKVNSTLEQYVFCTV-360  361-NALPETTADIVVFDEISMATNYDLSVVNAR-390  391-LRAK**HYVYI**GDPAQLPAPRTLLTKGTLEPE-420  421-**YFNSV**CRLMKTIGPDMFLGTCRRCPAEIVD-450  451-TVSALVYDNKLKAHKDKSAQCFKMFYK**GVI**-480  481-**TH**DVSSAINRPQI**GVVRE**FLTRNPAWRKAV-510  511-FISPYNSQNAVASKILGLPTQTVDSSQGSE-540  541-Y**DYVIF**TQTTETAHSCNVNRFNVAITRAKV-570  571-GILCIMSDRDLYDKLQFTSLEIPRRNVATL-600  601-Q | Total sites: 86 (14.30%)  Sites within APR: 9 (10.47%)  Sites outside APR: 77 (89.53%) |
| **NSP14** | 1-AENVTGLFKDCSKVITGLHPTQAPTHLSVD-30  31-TKFKTEG**LCVDI**PGIPKDMTYRRLISMMGF-60  61-KMNYQVNGYPNMFITREEAIRHVRAWIGFD-90  91-VEGCHATREAVGTNLPLQLGFST**GVNLV**AV-120  121-PTGYVDTPNNTDFSRVSAKPPPGDQFKHLI-150  151-PLMYKGLPW**NVVRI**KIVQMLSDTLKNLSDR-180  181-VVFVLWAHGFELTSMKYFVKIGP**ERTCCLC**-210  211-**D**RRATCFSTASDTYACWHHSI**GFDYV**YNPF-240  241-MIDVQQWGFTGNLQSNHDLYCQVHGNAHVA-270  271-SCDAIMTRCLAVHECFVKRVDWTIEYPIIG-300  301-DELKINAACRKVQHMVVKAALLADKFPVLH-330  331-DIGNPKAIKCVPQADVEWKFYDAQPCSDKA-360  361-YKIEELFYSYATHSDKFTD**GVCLF**WNCNVD-390  391-RYPANSIVCRFDTRVLSNLNLPGCDG**GSLY**-420  421-**V**NKHAFHTPAFDKSAFVNLKQLPFFYYSDS-450  451-PCESHGKQ**VVSDI**DYVPLKSATCITRCNLG-480  481-GAVCRHHANE**YRLYL**DAYNMMISAGFSLWV-510  511-YKQFDTYNLWNTFTRLQ-527 | Total sites: 99 (18.78%)  Sites within APR: 11 (11.11%)  Sites outside APR: 88 (88.89%) |
| **NSP15** | 1-SLENVAFNVVNKGHFDGQQGEV**PVSIIN**NT-30  31-VYTKVDGVDVELFENKTTLPVNVAFELWAK-60  61-RNIKPVPEVKILNNL**GVDIA**A**NTVIW**DYKR-90  91-DAPAHISTI**GVCSM**TDIAKKPTETICAP**LT**-120  121-**VFF**DGRVDGQVDLFRNAR**NGVLIT**EGSVKG-150  151-LQPSVGPKQASLN**GVTLIG**EAVKT**QFNYY**K-180  181-KVD**GVVQQ**LPETYFTQSRNLQEFKPRSQME-210  211-IDFLELAMDEFIERYKLEGYAFEHIVYGDF-240  241-SHSQLGGLHLLIGLAKRFKESPFELEDFIP-270  271-MDST**VKNYF**ITDAQTGSSKCVCSVIDLLLD-300  301-**DFVEIIK**SQDLSVVSKVVKVTIDYTEISFM-330  331-LWCKDGHVETFYPKLQ-346 | Total sites: 39 (11.27%)  Sites within APR: 9 (23.07%)  Sites outside APR: 30 (76.93%) |
| **NSP16** | 1-SSQAWQPGVAMPNLYKMQRMLLEKCD**LQNY**-30  31-**G**DSATLPKGIMMNVAKY**TQLCQ**YLNTLTLA-60  61-VPYNMRVIHFGAGSDKGVAPGTAVLRQWLP-90  91-TGTLLVDSDLNDFVSDAD**STLIG**DCATVHT-120  121-ANK**WDLIIS**DMYDPKTKNVTKENDSKEGFF-150  151-TYICGFIQQKLALGGSVAIKITEHSWNADL-180  181-YKLMGHFAWWTAFVTNVNASSSE**AFLIG**CN-210  211-YLGKPREQIDGYVM**HANYI**FWRNTNPIQLS-240  241-SYSLFDMSKFPLKLRGTAVMSLKEGQINDM-270  271-ILSLLSK**GRLII**RENNRVVISSDVLVNN-298 | Total sites: 44 (14.76%)  Sites within APR: 5 (11.36%)  Sites outside APR: 39 (88.64%) |

**Figure S1**

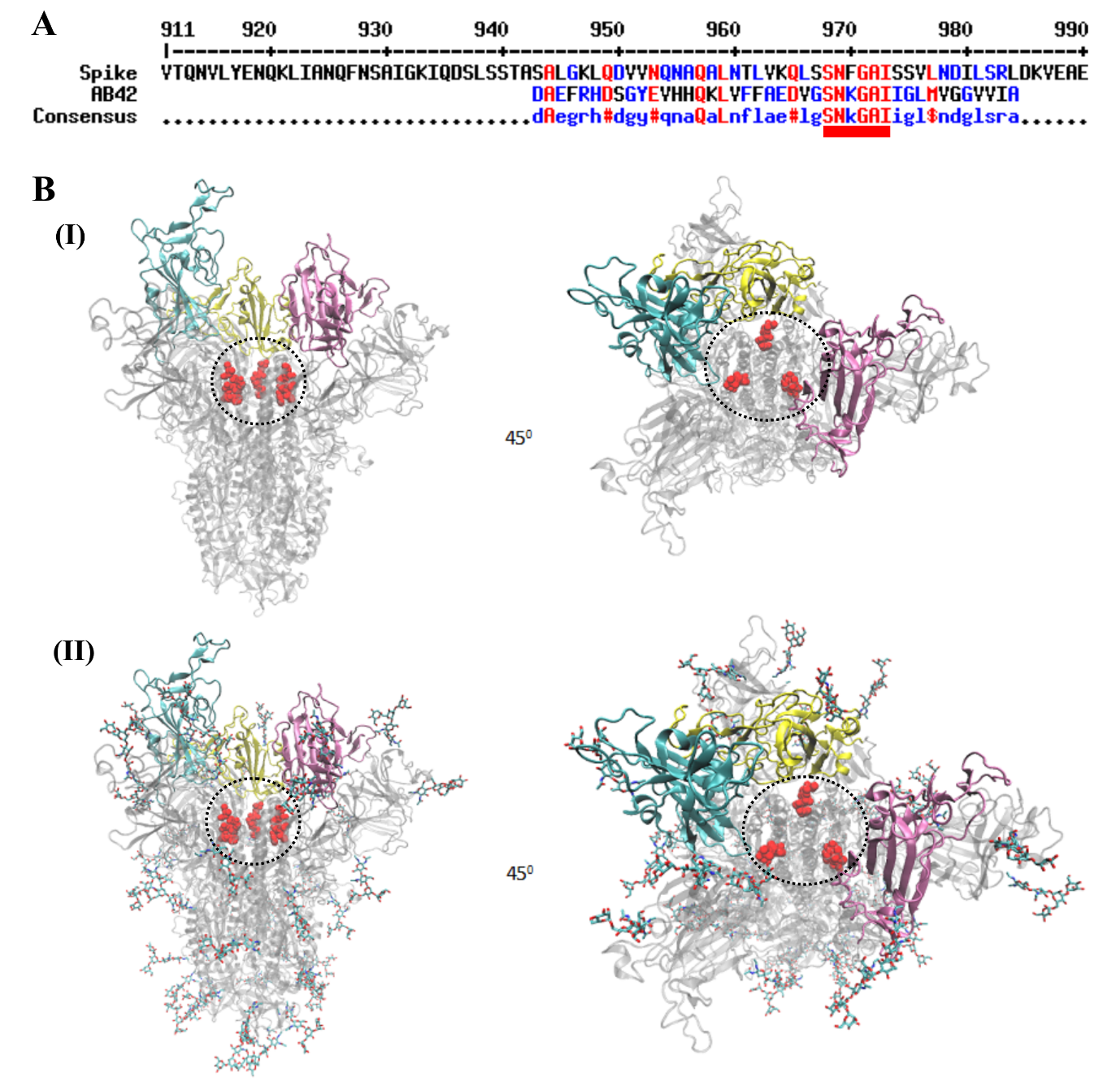

**Figure S1. Representation of amyloidogenic region in the open state conformation of spike protein.** (**A**) Multiple sequence alignment between SARS-CoV-2 spike and Aβ42. The aligned region of spike (residues 911 to 990) with Aβ42 is shown. The region _26_SNKGAI_31_ of Aβ42 is well aligned with _968_SNFGAI_973_ of spike. (**B**) Representation of spike in the open state. In cyan, purple, and yellow we represented the RBDs of each protomer and in red VDW the _968_SNFGAI_973_ sequence in the three protomers. (**I**) Spike without the glycan shield and (**II**) spike coated with glycans.

**Mass spectrometry data of crude peptides**

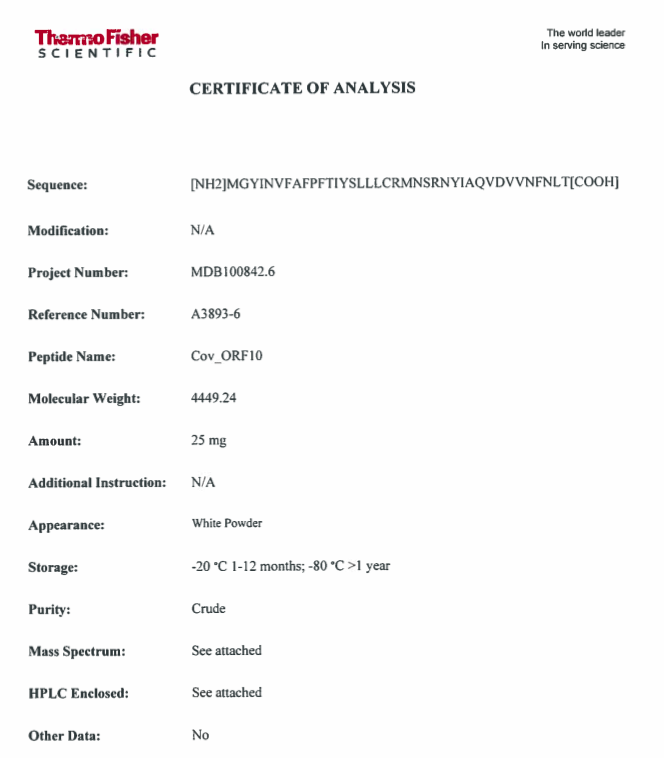

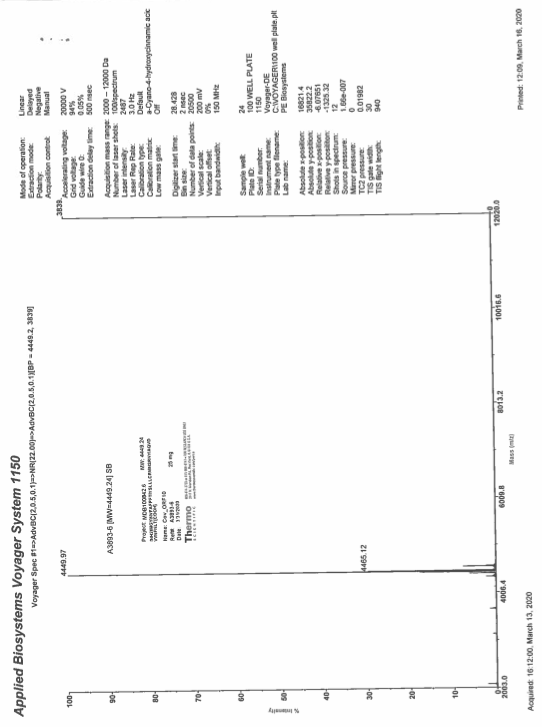
